# Genetic effects on *planum temporale* asymmetry and their limited relevance to neurodevelopmental disorders, intelligence or educational attainment

**DOI:** 10.1101/622381

**Authors:** Amaia Carrion-Castillo, Antonietta Pepe, Xiang-Zhen Kong, Simon E Fisher, Bernard Mazoyer, Nathalie Tzourio-Mazoyer, Fabrice Crivello, Clyde Francks

**Affiliations:** Language and Genetics Department, Max Planck Institute for Psycholinguistics, Nijmegen, The Netherlands; Groupe d’Imagerie Neurofonctionnelle, Institut des Maladies Neurodégénératives, Centre National de la Recherche Scientifique, Commissariat à l’Energie Atomique, et Université de Bordeaux, Bordeaux, France; Donders Institute for Brain, Cognition and Behaviour, Radboud University, Nijmegen, The Netherlands

**Author notes:** Corresponding author Clyde Francks, Max Planck Institute for Psycholinguistics, PO BOX 310, 6500 AH Nijmegen, The Netherlands.

**Keywords:** planum temporale, UK Biobank, genome-wide association, genetic correlation, neurodevelopmental disorders, brain asymmetry

## Abstract

Previous studies have suggested that altered asymmetry of the planum temporale (PT) is associated with neurodevelopmental disorders, including dyslexia, schizophrenia, and autism. Shared genetic factors have been suggested to link PT asymmetry to these disorders. In a dataset of unrelated subjects from the general population (UK Biobank, N= 18,057), we found that PT volume asymmetry had a significant heritability of roughly 14%. In genome-wide association analysis, two loci were significantly associated with PT asymmetry, including a coding polymorphism within the gene *ITIH5* that is predicted to affect the protein’s function and to be deleterious (rs41298373, P=2.01×10-15), and a locus that affects the expression of the genes *BOK* and *DTYMK* (rs7420166, P=7.54×10-10). *DTYMK* showed left-right asymmetry of mRNA expression in post mortem PT tissue. Cortex-wide mapping of these SNP effects revealed influences on asymmetry that went somewhat beyond the PT. Using publicly available genome-wide association statistics from large-scale studies, we saw no significant genetic correlations of PT asymmetry with autism spectrum disorder, attention deficit hyperactivity disorder, schizophrenia, educational attainment or intelligence. Of the top two individual loci associated with PT asymmetry, rs41298373 showed a tentative association with intelligence (unadjusted P=0.025), while the locus at *BOK/DTYMK* showed tentative association with educational attainment (unadjusted Ps<0.05). These findings provide novel insights into the genetic contributions to human brain asymmetry, but do not support a substantial polygenic association of PT asymmetry with cognitive variation and mental disorders, as far as can be discerned with current sample sizes.

## 1 Introduction

The *planum temporale* (PT) is a triangular-shaped region of the cerebral cortex, located posteriorly on the superior temporal gyrus (STG) (Altarelli, et al., 2014). Geschwind and Levitsky (1968) first described left-right asymmetry of PT surface area fifty years ago. They analyzed 100 post-mortem brains and observed a leftward asymmetry (left>right) in 65 of them, rightward asymmetry in 10, and approximate symmetry in 24. Re-analysis of the same brains emphasised that this variation is continuous in nature (Galaburda, Corsiglia, Rosen, & Sherman, 1987). Further studies have consistently shown that the PT is leftward lateralized in the general population, and that the left PT can be up to 50% larger than the right, although the precise anatomical definition affects this estimate (Toga & Thompson, 2003; Watkins, et al., 2001; Tzourio-Mazoyer, et al., 2010; Tzourio-Mazoyer & Mazoyer, 2017; Tzourio-Mazoyer, et al., 2018).

Lesion and functional studies suggest that the PT is implicated in a number of language-related processes, including auditory and phonological processing, and language comprehension (Dronkers, Wilkins, Van Valin, Redfern, & Jaeger, 2004). Initial reports proposed that PT asymmetry may be an anatomical marker for left-hemisphere dominance for language processing (Geschwind & Levitsky, 1968; Foundas, Leonard, Gilmore, Fennell, & Heilman, 1994). However, recent studies have found that there is no correlation between PT asymmetry and functional language lateralization as assessed at the whole hemisphere level (Keller, et al., 2011; Tzourio-Mazoyer, Crivello, & Mazoyer, 2017). Rather, it seems that PT anatomical asymmetry is associated with functional lateralization only locally, in directly neighbouring auditory areas (Tzourio-Mazoyer, Crivello, & Mazoyer, 2017).

Alterations of the typical form of PT asymmetry have been reported in dyslexia. In a post-mortem analysis, Galaburda et al. (Galaburda, Sherman, Rosen, Aboitiz, & Geschwind, 1985) observed symmetrical *plana temporale* in the brains of all dyslexic individuals that they studied (4 males and 3 females), but subsequent studies using neuroimaging have found contradictory results (see Table 1 in Altarelli et al. (2014) for an overview). Using magnetic resonance imaging (MRI), Altarelli et al. (2014) manually outlined the PT according to the same anatomical criteria as in earlier post mortem studies, in 35 dyslexic cases and 46 controls. They found an increased proportion of rightward surface area asymmetry among dyslexic boys specifically, and suggested that previous mixed results may have arisen due to different anatomical definitions of the PT, and not accounting adequately for sex.

**Table 1:**
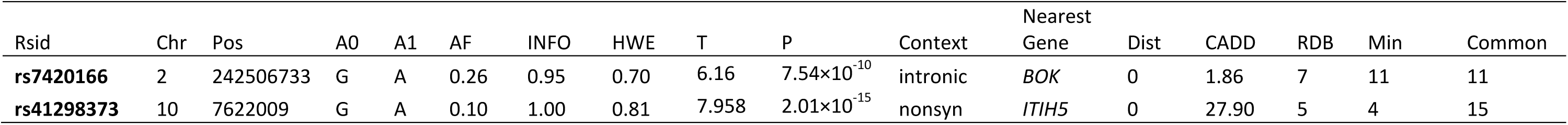
Significantly associated loci with PT asymmetry in the UK Biobank dataset. Rsid: Rs identity of the lead SNP within the locus. Pos: chromosome position of the SNP in the hg19 human reference genome. A0 and A1 are the SNP alleles. A1 is the ‘effect allele’, i.e. the direction of the association T statistic with the PT AI indicates the additive effect of each A1 allele. AF: allele frequency of the effect allele. INFO: imputation quality score. HWE: Hardy-Weinberg p-value. T: association T-statistic. P: association P value. Context: location of the SNP with respect to the nearest gene, and consequence at the protein level if exonic (nonsyn: non synonymous). Dist: distance to the nearest gene in basepairs. CADD: Combined Annotation Dependent Depletion (CADD) score (scores greater than 15 indicate deleterious SNPs). RDB: regulome DB categorical score (from 1a to 7). 1a is the highest score for SNPs with the most biological evidence to be a regulatory element. Chrom min and chrom common: the minimum and most common chromatin states across 127 tissue/cell types. States are described using a 15 point scale, with the lower the chromatin score the greater the accessibility to the genome at this site (scores of less than 8 indicate an open chromatin region)

| Rsid | Chr | Pos | A0 | A1 | AF | INFO | HWE | T | P | Context | Nearest Gene | Dist | CADD | RDB | Min | Common |
| --- | --- | --- | --- | --- | --- | --- | --- | --- | --- | --- | --- | --- | --- | --- | --- | --- |
| **rs7420166** | 2 | 242506733 | G | A | 0.26 | 0.95 | 0.70 | 6.16 | $7.54 \times 10^{-10}$ | intronic | *BOK* | 0 | 1.86 | 7 | 11 | 11 |
| **rs41298373** | 10 | 7622009 | G | A | 0.10 | 1.00 | 0.81 | 7.958 | $2.01 \times 10^{-15}$ | nonsyn | *ITIH5* | 0 | 27.90 | 5 | 4 | 15 |

A reduction of PT grey matter volume asymmetry has also been reported in people with autism spectrum disorder (ASD), with a smaller left PT in cases causing a change of asymmetry (Rojas, Bawn, Benkers, Reite, & Rogers, 2002; Rojas, Camou, Reite, & Rogers, 2005). A reduction of the left PT has been proposed to relate to delays in language acquisition, which can occur in some ASD patients (Knaus, Kamps, Foundas, & Tager-Flusberg, 2017). Consistent with this, a recent study of brain regional growth rates in newborns with congenital heart diseases reported a strong correlation between the growth rate of left PT and language scores at 12 months of age (Jakab, et al., 2019). Furthermore, a meta-analysis of ten studies concluded that a reduction of PT area asymmetry is a feature of schizophrenia (Sommer, Ramsey, Kahn, Aleman, & Bouma, 2001), although it is not clear whether this change in asymmetry is driven by a reduced left PT or increased right PT in cases, or both (Shapleske, Rossell, Woodruff, & David, 1999; Hirayasu, et al., 2000; Hasan, et al., 2011). A reduction of PT asymmetry in terms of area and thickness measures has also been observed in schizophrenia (SCZ) (Ratnanather, et al., 2013). Auditory hallucinations can occur in SCZ (Morch-Johnsen, et al., 2017), which may conceivably relate to PT functions as secondary auditory cortex (Crow, Ball, Bloom, & al, 1989).

PT asymmetry is established early in development, as evidenced by studies of fetuses (Chi, Dooling, & Gilles, 1977), preterm newborns (Dubois, et al., 2010) and infants (Wada, Clarke, & Hamm, 1975; Li, et al., 2014), such that a genetic-developmental program is likely to underlie this asymmetry. Shared genetic contributions to schizophrenia and PT asymmetry have long been postulated (Sommer, Ramsey, Kahn, Aleman, & Bouma, 2001; 1993), although early hypotheses were formulated around single genetic loci of large effect, and it is now undisputed that there is a substantial polygenic contribution to schizophrenia, with significant SNP-based heritability and multiple susceptibility loci identified (Ripke, et al., 2014). A shared genetic contribution to PT asymmetry and dyslexia has also been suggested, on the basis that pre-readers and adolescents at familial risk of dyslexia have shown a reduced leftward PT asymmetry compared to those with no familial risk (Vanderauwera, et al., 2018).

A previous genome-wide association scan (GWAS) meta-analysis for PT volume asymmetry, in just over 3000 healthy adult subjects, did not find individual loci which were significant after correcting for multiple testing over the whole genome (Guadalupe, et al., 2015). However, a significant enrichment of association signals was found within sets of genes with roles in steroid hormone receptor activity and steroid metabolic processes. Gene sets involved in steroid hormone biology were tested because PT asymmetry showed the most significant sex difference of all cortical regional asymmetries, with males more leftward asymmetric on average (Guadalupe, et al., 2015). It has also been reported that foetal testosterone levels relate to the development of grey matter asymmetries of some cortical regions, including the PT (Lombardo, et al., 2012). In fact, an extensive and influential theory was developed by Geschwind and Galaburda which involves sex hormones, brain laterality and disorder susceptibility (Geschwind & Galaburda, 1985), which still motivates current research, although various elements remain unproven (Hollier, Maybery, Keelan, Hickey, & Whitehouse, 2014; Papadatou-Pastou & Martin, 2017).

In the present study, we used the UK Biobank MRI dataset (N=18,057) to perform a more highly powered genetic analysis of PT asymmetry than has previously been possible. This sample size also permitted, for the first time, the estimation of SNP-based heritability of PT asymmetry, and its genetic correlations with other traits. Our goals were (i) to identify individual genetic loci and gene sets related to variation in PT asymmetry, which can provide novel insights into genetic contributions to brain laterality, and (ii) to examine whether polygenic influences on PT asymmetry also impact cognitive traits and psychiatric disorders. For the latter purpose we made use of summary statistics from large-scale GWAS studies of ASD and schizophrenia. There are currently no large-scale GWAS results available for dyslexia (based on sample sizes of over 10,000), but we also used GWAS summary statistics for intelligence and educational attainment (EA), as these traits are correlated both phenotypically and genetically with language and reading abilities in the general population (Verhoef, et al., 2019). Finally we included analysis of GWAS summary statistics for attention deficit/hyperactivity disorder (ADHD), as this disorder is often comorbid with dyslexia and negatively genetically correlated with reading measures (Verhoef, et al., 2019).

## 2 Materials and methods

### 2.1 Dataset

Data were obtained from the UK Biobank cohort, as part of research application 16066, with Clyde Francks as the principal applicant. This is a general adult population cohort. The data collection in the UK Biobank, including the consent procedure, has been described elsewhere (Sudlow, et al., 2015). Informed consent was obtained by the UK Biobank for all participants. For this study we used data from the October 2018 release of 22,392 participants’ brain MRI data, together with the genome-wide genetic data from the same participants. The age range of these participants was from 44 to 80 years (median 63.9), and 11,269 were female, 10,551 were male.

### 2.2 Genetic quality control

We excluded subjects with a mismatch of their self-reported and genetically inferred sex, with putative aneuploidies, or who were outliers based on heterozygosity (PC corrected heterozygosity > 0.19) and genotype missingness (missing rate > 0.05) (Bycroft, et al., 2018). We further restricted the analysis to participants with ‘British ancestry’, as defined by Bycroft et al (‘in.white.British.ancestry.subset’) (Bycroft, et al., 2018). We randomly excluded one subject from each pair with a kinship coefficient > 0.0442, as defined within the UK Biobank relatedness file ‘*ukb1606_rel_s488366.dat*’, unless one of the pair had no PT L and R measures (see below), in which case that subject was selected for exclusion. This resulted in 18,057 subjects passing genetic QC.

### 2.3 PT volume and asymmetry measures

The UK Biobank provided various derived brain imaging measures (see Miller et al. (Miller, et al., 2016) and documentation, https://biobank.ctsu.ox.ac.uk/crystal/docs/brain_mri.pdf). Briefly, tissue-type segmentation (Zhang, Brady, & Smith, 2001) was performed from T1 images using FSL FAST (Smith, et al., 2004; Jenkinson, Beckmann, Behrens, Woolrich, & Smith, 2012), and 44 cerebral cortical regions were defined per hemisphere based on the Harvard-Oxford probability atlas (distributed with the FSL software package; http://fsl.fmrib.ox.ac.uk/fsl/). For the present study we focus on the PT as defined in the Harvard-Oxford atlas, which makes use of an anatomical definition close to that used in the original post mortem studies of this region (see Introduction). Grey matter volumes of the left and right PT were calculated per subject by summing the voxel-wise grey matter volume estimates within each region.

We extracted the left (L) and right (R) PT grey matter volumes (UKB field IDs= f.25872.2.0 and f.25873.2.0), and calculated an asymmetry index (AI) per individual subject as (L-R)/((L+R)/2). Given this definition, a positive AI reflects a leftward asymmetry of PT grey matter volume. The main focus of the present study was on the AI, but the L and R measures separately, as well as total brain volume (TBV, UKB field ID: 25010, ‘Volume of brain, grey+white matter’) were also analyzed in some contexts, to aid the interpretation of findings involving the AI. All analyses, data transformations and filtering steps were done in R (version 3.3.2).

Outliers were removed at greater than 4 standard deviations from the mean (calculated after the exclusion of some subjects on the basis of genetic quality control, see below) separately for each trait (L, R, AI, and TBV). For each trait we then performed linear regression, using the *lm* function in R, and extracted the residuals after correcting for sex, age, nonlinear age (zage2), genotyping array, scanner position parameters (x=lateral, y=transversal, z=longitudinal), and the first ten principal components (PCs) which capture genome-wide population structure in the genotype data and were made available by the UK Biobank (Bycroft, et al., 2018) (Table S1). The distribution of each trait was visually inspected before and after residualizing (Figure S1). The outlier exclusion, together with some missing PT data in the UK Biobank release, and some missing or outlier subjects for some covariates, meant that between 18,037 and 18,053 subjects were available with both post-QC genetic data and PT or TBV measures, depending on the specific measure (Table S2).

Initially we also considered self-reported handedness as a potential covariate effect (UK Biobank field ID f.1707.0.0, coded as right-handed vs left-handed, and excluding ambilaterals), but there was no association with PT asymmetry (t=-0.62, p-value-0.53), and we therefore dropped handedness from the analyses.

### 2.4 Heritability and genetic correlations between PT measures and total brain volume

550,192 autosomal, directly genotyped single nucleotide polymorphisms (SNPs) with minor allele frequencies (MAF) > 0.01, genotyping rate >0.95 and Hardy-Weinberg equilibrium (HWE) p>1×10^−6^ were used to build a genetic relationship matrix (GRM) using GCTA (version 1.26.0) (Yang, et al., 2010). For the specific purposes of heritability and genetic correlation analysis, we further excluded samples with a genotyping rate < 98% and a kinship coefficient higher than 0.025 based in this GRM (N=17,221), resulting in sample sizes of between 17,185 and 17,209 for these particular analyses, depending on the measure or pair of measures (Table S3). Genome-based restricted maximum likelihood (GREML) (Yang, et al., 2010) analyses were performed to estimate the SNP-based heritabilities (h^2^_SNP_) of the AI, L, R, and TBV measures. Bivariate GREML was used to estimate genetic correlations between measures.

### 2.5 Genome wide association scanning and annotation

Imputed SNP genotype data (bgen files; imputed data v3 -released March 2018) were extracted for the samples with PT data (N=18,057), and snp-level statistics were then computed within this set using QCtools (v.2.0.1). GWAS was performed separately for each of the residualized PT AI, L and R measures, using imputed genotype dosages and an additive model, with BGENIE (v1.2) (Bycroft, et al., 2018). Sample sizes ranged from 18,037 to 18,053 depending on the specific measure (Table S2). We excluded SNPs with minor allele frequencies below 0.001, Hardy-Weinberg p-value below 1×10^−7^ or imputation quality INFO scores below 0.7 (the latter as provided by the UKBiobank with the imputed data), which resulted in 15,120,452 analyzed SNPs in the GWAS. Manhattan plots and QQplots were made using the “qqman” R (Turner, 2017) package, and regional association plots were made using LocusZoom v1.4 (Pruim, et al., 2010).

The summary statistics from the GWAS for PT AI were loaded into the annotation software FUMA (Watanabe, Taskesen, van Bochoven, & Posthuma, 2017). Significantly associated SNPs were considered as those with pointwise P < 5 × 10^-8^, to correct for multiple testing over the whole genome. These SNPs were matched based on chromosome, base pair position, reference and non-reference alleles to a database containing variant level functional annotations: ANNOVAR (Wang, Li, & Hakonarson, 2010), which compiles information including Combined Annotation Dependent Depletion (CADD) scores (v1.3) (Kircher, et al., 2014), Regulome DB (RDB) scores (v1.1) (Boyle, et al., 2012) and chromatin states (Dunham, et al., 2012; Kundaje, et al., 2015; Ernst & Kellis, 2012) (see Results).

### 2.6 Gene-set enrichment analysis

Gene-set enrichment analysis was applied to the GWAS results for the PT AI, using the software MAGMA v1.06b (de Leeuw, Mooij, Heskes, & Posthuma, 2015). The basis of this analysis is to first assign a single association score to each gene based on the GWAS data, and then test whether the genes in a given set show, on average, more evidence for association with the trait in question than the rest of the genes in the genome, for which scores could be calculated (hence this is sometimes called ‘competitive’ analysis). All SNPs that were located within protein coding genes, and up to 35kb upstream and 10kb downstream from them, were used to calculate gene scores within MAGMA using the sum of −log(SNP p-value)(gene-model: ‘snp-wise=mean’). Gene locations and boundary definitions were used from the NCBI build 37, and linkage disequilibrium was controlled using the 1000 genomes phase 3 release (ftp://ftp.1000genomes.ebi.ac.uk/vol1/ftp/release/20130502/) provided by MAGMA (de Leeuw, Mooij, Heskes, & Posthuma, 2015).We tested for enrichment according to the Gene Ontology (GO) classification system (Ashburner, et al., 2000; Consortium G., 2017), in which sets are defined according to molecular functions, cellular components or biological processes. GO terms were obtained from Msigdb v6.1 (http://software.broadinstitute.org/gsea/msigdb/collections.jsp#C5). A total of 5,875 GO sets (with a minimum of 10 genes per set) were tested, and Bonferroni multiple testing correction was applied to obtain corrected p-values.

We also interrogated two specific GO gene-sets related to steroid hormone biology (‘steroid hormone receptor activity’ and ‘steroid metabolic process’) which were enriched for association signals with PT asymmetry in a previous, smaller GWAS study of PT asymmetry (see Introduction) (Guadalupe, et al., 2015).

### 2.7 Further analysis of rs41298373 and rs7420166

Two SNPs, rs41298373 and rs7420166, were subject to further characterisation and analysis because they were significantly associated with the PT AI in the GWAS. Rs41298373 is a protein coding variant with a predicted deleterious impact, located in the gene *ITIH5*, while rs7420166 was the lead SNP at the chromosome 2q37 locus, located within an intron of the gene *BOK* (see Results).

#### 2.7.1 Bioinformatic characterisation

Evolutionary conservation was assessed using Genomic Evolutionary Rate Profiling (GERP), which provides position-specific estimates of evolutionary constraint (Davydov, et al., 2010). Rs41298373, as a coding SNP, was also annotated with SIFT (Kumar, Henikoff, & Ng, 2009) and Polyphen-2 (Adzhubei, Jordan, & Sunyaev, 2013). Polyphen-2 uses aminoacid conservation, structure, and protein-level annotation, while SIFT uses aminoacid sequence homology to infer possible impacts of amino acid substitutions on protein structure. FUMA (Watanabe, Taskesen, van Bochoven, & Posthuma, 2017) was used to query expression QTL (eQTL) effects of rs7420166 and its proxy SNPs (LD r^2^>0.8). We queried the largest available eQTL datasets, which are based on whole blood (Westra, et al., 2013; Zhernakova, et al., 2017), as well as eQTL data derived from smaller samples but using adult human brain tissue: cortex (GTEx v7) (Consortium G., 2015), temporal cortex (Braineac) (Ramasamy, et al., 2014), dorsolateral cortex (xQTL) (Ng, et al., 2017). These analyses would implicate BOK but also another nearby gene, DTYMK (see Results).

The spatio-temporal expression patterns of *ITIH5, BOK*, and *DTYMK* were characterized by querying data from several sources. For human adult expression data: GTEx Consortium (Consortium G., 2015), Allen Human Brain Atlas (Hawrylycz, et al., 2012), BRAINEAC (Ramasamy, et al., 2014). For human developmental data: the BrainSpan database (Miller, et al., 2014). For human adult single cell RNA sequencing (middle temporal gyrus): Allen Brain Atlas (http://celltypes.brain-map.org/rnaseq/human). For transcriptome profiles of brain cell-types (glia, neurons, and vascular cells): the ‘brainrnaseq’ database (http://www.brainrnaseq.org/) from human adult samples (temporal lobe cortex) (Zhang, et al., 2016) and mouse (Zhang, et al., 2014).

Using the Allen Brain Atlas microarray data, which are based on six post mortem brains (only two have data from both hemispheres), we also assessed whether the expression of *ITIH5, BOK* or *DTYMK* in the PT differed between hemispheres, by modelling expression as a function of probe and hemisphere, and including subject as a random factor (expression ∼ (1|subject) + probe + hemisphere), using the *lmer* function in the ‘lme4’ package in R. We used all data assigned to the region ‘PLT’ as defined in the Allen data.

#### 2.7.2 Brain-wide effects on grey matter asymmetry

We performed grey matter voxel-based-morphometry (VBM) association analyses for each of the SNPs rs41298373 and rs7420166, to assess their potential effects on asymmetry brain-wide. Briefly, voxel-wise grey matter volume was assessed with FSLVBM (Douaud, et al., 2007)(http://fsl.fmrib.ox.ac.uk/fsl/fslwiki/FSLVBM) carried out with FSL (v5.0) tools (Smith, et al., 2004; Jenkinson, Beckmann, Behrens, Woolrich, & Smith, 2012). Subject’s grey matter images were non-linearly normalized onto a left-right-symmetric grey matter template in the standard MNI 152 space. The generated maps were smoothed with isotropic Gaussian kernel (sigma=2.55, full-width at half-maximum=6mm). Next, voxel-wise asymmetry index (AI) maps were calculated using the formula (L-R)/((L+R)/2) for each pair of left-right corresponding voxels, per subject. Finally, we ran association analyses of rs41298373 and rs7420166 with each voxel AI using mri_glmfit (http://surfer.nmr.mgh.harvard.edu/fswiki/mri_glmfit) from FreeSurfer (v6.0) (Dale, Fischl, & Sereno, 1999; Fischl, Sereno, & Dale, 1999), including the covariates listed in Table S1, and testing the additive effect of each genotype (hard-called genotypes, call probability threshold 0.9, resulting in N=18052 subjects for rs41298373 and N=16663 for rs7420166). Multiple comparison correction was performed on statistical maps using the 3dClustSim program implemented in AFNI (version 16.2.19). Thresholds of voxel-level p <0.001 (Z=3.29) and cluster-level p <0.01 (cluster size >=40 voxels;) were used based on Monte Carlo simulation in the left hemisphere mask. Results were mapped to left-hemisphere surface space for visualization of cerebral cortical effects, using BrainNet Viewer (v1.63) (Xia, Wang, & He, 2013) (the ‘extremum voxel’ option).

#### 2.7.3 Vertex-wise analysis of rs41298373 and rs7420166

We performed a surface-based analysis in a subset of 8,778 subjects (from an earlier release of UK Biobank data) for which cortical surface extraction was obtained using FreeSurfer (v6.0) (Dale, Fischl, & Sereno, 1999; Fischl, Sereno, & Dale, 1999) (https://surfer.nmr.mgh.harvard.edu/) with 1 mm^3^ T1 and FLAIR acquisitions as inputs. A quality control procedure (ENIGMA Consortium protocol, http://enigma.ini.usc.edu) for FreeSurfer surface reconstruction was performed on the cortical surface area and cortical thickness estimates at both hemispheric and regional (Desikan, et al., 2006) levels. This involved producing interactive web-page reports for group-level statistics of the hemispheric and regional measures, and subject-specific web-page reports for surface reconstruction accuracy. Visual inspection of the subject-specific reports was performed on individuals flagged as possible outliers (if they deviated more than 4 standard deviations from the mean) based on the group-level statistics of any of the aforementioned measures, and resulted in the exclusion of 189 subjects. The left and right cortical surfaces of the 8589 remaining participants were then warped into a left/right symmetrical standard space (*fsaverage_sym*) using a surface-based inter-hemispheric co-registration procedure acting at the vertex level (Maingault, Tzourio-Mazoyer, Mazoyer, & Crivello, 2016; Greve, et al., 2013). This inter-hemispheric alignment step was done twice: once on the left hemisphere of the symmetrical *fsaverage_sym* template, and once on its right hemisphere counterpart. In these two surface models, and for each participant, vertex-level cortical thickness and cortical surface area left-right difference (L-R) maps were generated. The two cortical thickness asymmetry maps (one for each hemisphere of *fsaverage_sym surface model*) were further averaged to produce a more robust left-right difference map, which was unbiased for positional structural asymmetries between hemispheres. The same was done for cortical surface area asymmetry. Again, group-based quality control of the inter-hemispheric vertex-wise alignment was performed, which led to a usable sample of 8578 subjects (11 additional subjects excluded for failure of inter-hemispheric registration). Next, the cortical thickness and surface area vertex-wise left-right difference maps were smoothed with a 15 mm^2^ FWHM surface-based Gaussian kernel, and processed with the previously mentioned *mri_glmfit* Freesurfer command-line to search for possible associations of rs41298373 and rs7420166 with each vertex, using the same statistical model as in the VBM analysis (above), and covariates listed in Table S1. Coupled with the previous genetic quality control (see above) and the available demographic data, and testing the additive effect of each genotype (hard-called genotypes, call probability threshold 0.9), there were N=7231 subjects for rs41298373 and N=6685 for rs7420166. Association maps with the two SNPs were obtained after cluster wise correction at p<0.05 for multiple comparisons, by parametric Gaussian based Monte Carlo simulations, and 0.005 as the cluster forming threshold.

### 2.8 Disorders, cognitive and behavioural traits

#### 2.8.1 Genome-wide genetic correlation

We estimated genetic correlations of the PT AI, L and R measures with SCZ, ASD, ADHD, intelligence, and EA. This analysis was based on GWAS summary statistics for the PT measures as generated in this study, together with publicly available GWAS summary statistics for ASD (Grove, et al., 2017), SCZ (Ripke, et al., 2014), ADHD (Demontis, et al., 2018), intelligence (Savage, et al., 2018) and educational attainment (EA) (Lee, et al., 2018). Sample sizes used for each of these GWAS, prevalence and other parameters are summarized in Table S7; we used the publicly available summary statistics of the largest-to-date GWASes for each of these traits.

To account for different possible genetic models, we assessed genetic correlation using two different methods which are both designed to make use of summary statistics, i.e. do not require individual level data: LD score regression (LDSC, v.1.0.0) (Bulik-Sullivan, et al., 2015) and SumHer (Speed & Balding, 2018). For LDSC, pre-computed LD scores based on 1000 Genomes European-descent data (https://data.broadinstitute.org/alkesgroup/LDSCORE/) were used to model LD. We only used summary statistics for SNPs (MAF> 0.01 and INFO>0.9) that were shared with the reference panel. For SumHer (as implemented in ldak v.5.0), the same pre-computed LD scores were used, and two different models were run: the ldak-model in which the expected contribution of each SNP to heritability is weighted by its frequency (*–weights sumsect/weights.short –power −0.25*) and the gcta-model, in which all SNPs are expected to contribute equally (*–ignore-weights YES –power −1*).

We estimated the minimum power to detect genetic correlations across these pairs of traits and datasets at alpha <0.05 using the GCTA-GREML power calculator (Visscher, et al., 2014), given the sample sizes involved, and according to SNP-based heritability estimates obtained from LDSC and SumHer (see Figure S3). Tables S3 and S7 summarize all the parameters per trait that went into the power calculations. Since LD score regression (Bulik-Sullivan, et al., 2015) and SumHer (Speed & Balding, 2018) utilize summary statistics while GCTA relies on the individual genotype data, the true power is likely to be slightly lower for the former two; however, the GCTA-GREML power calculator gives an indicative estimate.

#### 2.8.2 Top SNPs and gene sets associated with PT asymmetry

For three individually significant SNPs from the GWAS of PT AI in the UK Biobank (see Results), we looked up these SNPs in the publicly available GWAS results for ASD, SCZ, ADHD, intelligence and EA. In case the SNP was not present in the publicly available GWAS results, we identified proxy SNPs in high LD (r^2^>0.8) with them, as calculated within the UKB imaging dataset using plink (v1.90) (see Table S9).

## 3 Results

Consistent with a previous report (Guadalupe, et al., 2015) based on roughly 3000 subjects from the Netherlands and Germany, the PT grey matter volume in the UK Biobank dataset of 18,049 healthy adult subjects, as measured using the Harvard-Oxford atlas, was leftward asymmetrical at the population level, with males having a slightly more pronounced leftward asymmetry than females (t= −11.38, p-value= 6.17×10^−30^, Figure S2).

### 3.1 Genetic architecture of PT asymmetry

The PT AI showed a significant SNP-based heritability, as assessed with GCTA (Yang, et al., 2010) (h^2^_SNP_(AI)= 0.139 se=0.035, p=1.38×10^−5^)). The SNP-based heritabilites for the left and right PT volumes, considered separately, were also significant and similar in magnitude to each other (h^2^_SNP_(left)= 0.454, h^2^_SNP_(right)= 0.414). However, the genetic correlation between left and right PT volumes was significantly different from 1 (ρ_(L,R)_= 0.85, p=0.009, null hypothesis = different to 1), which indicates that there are some genetic effects which are not shared between the left and right PT, and therefore constitute heritable contributions to PT asymmetry. There was a low but significant phenotypic correlation between AI and TBV (r=0.071, p=9.40×10^−22^), and also a borderline significant genetic correlation between them (ρ=0.133, p=0.053, null hypothesis = different to 0)(Table S4). Therefore we did not adjust for TBV in the main genetic analysis of the PT AI, as we wished to capture all genetic influences on the PT AI, regardless of whether they might be shared with other aspects of brain anatomy (Aschard, Vilhjalmsson, Joshi, Price, & Kraft, 2015).

### 3.2 Genome wide association analysis of PT measures

Two loci on different chromosomes were significantly associated with the PT AI at a threshold p<5×10^−8^ (Table 1):

The most significant association with the PT AI was on chromosome 10p14 (rs41298373, p=2.01×10^−15^, Figure 1b-S4). The minor A allele of rs41298373 (MAF=0.10) was associated with an increased leftward asymmetry of PT volume (T=6.18, Figure S5, Table S5). In fact this SNP was associated only with left-sided PT volume (p=6.48×10^−10^) and not with right-sided PT volume (p=0.13) (Figure S6, Table S5). Post hoc analysis showed that the association of rs41298373 with the PT AI was similar in males (beta=0.023, se=0.0047, p=7.76×10^−07^) and females (beta=0.028, se=0.0046, p=4.46×10^−10^), and the sex*rs41298373 interaction was not significant (F=0.541, p=0.462). There are no other SNPs within 500kb of rs41298373 that have LD with it at r^2^ > 0.5, which explains why this association signal appears relatively specific to rs41298373 (Figure S7). Nonetheless rs41298373 was directly genotyped and was in Hardy-Weinberg equilibrium (p=0.81), so that this result appears reliable.

**Figure 1.**
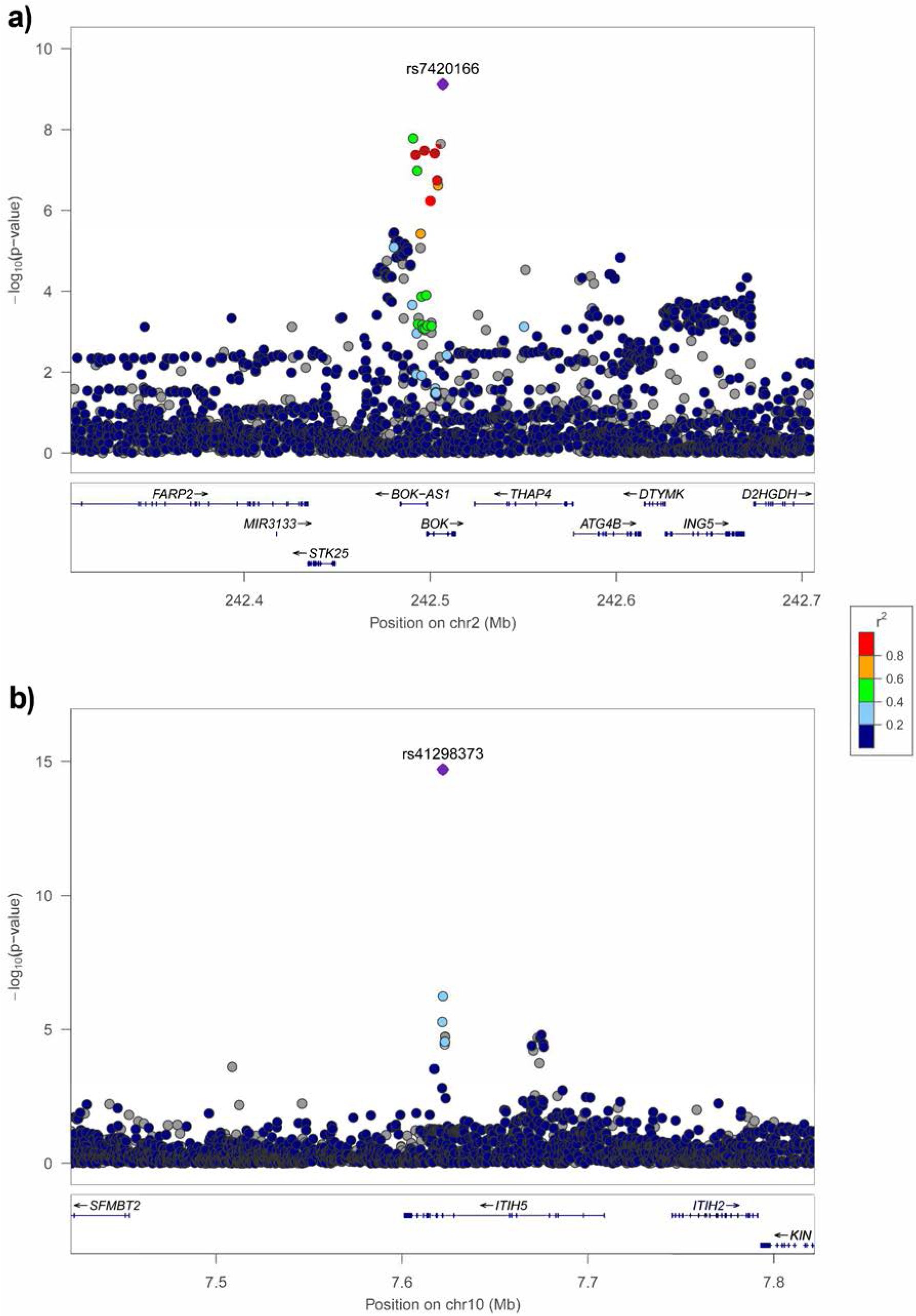
Regional association plot of part of **(a)** chr2q37 and **(b)** chr10p14, with PT volume asymmetry. The lead SNP (most significantly associated) is highlighted in violet, and the rest of the SNPs are coloured on the basis of their LD with the lead SNP (LD r^2^, within the 1KG EUR population).

The second most significant association with the PT AI was on 2q37, p=7.54×10^−10^ for the lead SNP rs7420166. There were 6 SNPs at this locus that had p values below 5×10^−8^, all in high-LD with each other (r^2^> 0.7 with the lead SNP) (Figure 1a). The effect of rs7420166 on PT asymmetry was driven by opposite effects on the left PT (T=3.70, p=2×10^−4^) and the right PT (T=-1.95, p=0.051), with the minor allele A (MAF = 0.256) resulting in increased leftward asymmetry (Table S5). Post hoc analysis showed that the association of rs7420166 with the PT AI was similar in males (beta=0.016, se=0.0034, p=4.12×10^−6^) and females (beta=0.0129, se=0.0031, p=4.47e-05), and the sex*rs41298373 interaction was not significant (F=1.58, p=0.209).

Neither of the lead SNPs rs41298373 and rs7420166, nor proxy SNPs in LD r^2^>0.8 with them, were included in the previously published GWAS meta-analysis of PT asymmetry in roughly 3000 subjects (Guadalupe, et al., 2015)(see Introduction).

No other loci surpassed the threshold p<5×10^−8^ in the GWAS of left PT, while a locus on chromosome 16q24.1 (lead SNP rs12932673, MAF=0.189, minor allele=G) just surpassed this threshold in the GWAS of the right PT volume (T=-5.59, p=2.315×10^−8^, Figure S6, Table S5). This SNP was also associated with the left PT volume to a lesser extent than the right (T=-2.76, p=5.845×10^−3^) such that the association with the PT AI was not even nominally significant (T=1.94, p=0.052).

### 3.3 Further characterisation of loci associated with PT asymmetry

Rs41298373 on 10p14 is an exonic nonsynonymous SNP within exon 9 of *ITIH5* (Inter-Alpha-Trypsin Inhibitor Heavy Chain Family Member 5)(transcript=ENST00000397145). The minor allele results in a substitution from alanine to valine (p.A376V), within the von Willebrand factor type A (VWA) domain of the protein (pfam=PF13768). The SNP is predicted to affect protein function by several bioinformatic tools (Polyphen = probably damaging, SIFT = deleterious, CADD = 27.9). The Genomic Evolutionary Rate Profiling (GERP) estimate for this genomic position is high (5.23), reflecting that the ancestral allele G is conserved across mammals. G is also the allele at this site in the reference genomes of archaic hominins (two Neanderthals and one Denisovan) (Green, et al., 2010; Meyer, et al., 2012). The minor allele A has a frequency around 10 % in most human populations, but is lower in African (2%) and East-Asian (0.7%) groups according to the gnomAD database (Lek, et al., 2016).

*ITIH5* encodes a heavy chain component of one of the inter-alpha-trypsin inhibitor (ITI) family members (Himmelfarb, et al., 2004). The ITI family contains multiple proteins (ITIP) that are made up of a light chain (bikunin) and a variable number of heavy chains (ITIH1-ITIH6), and are plasma serine protease inhibitors. They are involved in extracellular matrix stabilization and in prevention of tumor metastasis. ITIH5 is a known tumor suppressor gene, and it has been shown to be downregulated in breast cancer (Himmelfarb, et al., 2004).

The expression pattern of *ITIH5* is broad, being most highly expressed in adipose, artery, colon and mammary tissue (Consortium G., 2015). It also shows moderate expression in the brain throughout development (mean fragments per kilobase of exon per million reads mapped (FPKM) across all timepoints and regions in Brainspan = 2.86, range: 0.35-24.67; GTEx range of median Transcripts per million (TPM) across brain regions: 3.8-9.9), with a possible peak around the time of birth Figure S8, but with no particular spatial patterns discernible (BRAINEAC, GTEx, AllenBrainAtlas) (Ramasamy, et al., 2014; Consortium G., 2015; Hawrylycz, et al., 2012). In the Allen data from adult post mortem brain, the expression pattern within the PT did not reveal significant hemispheric differences, although there was a non-significant trend toward lower expression on the right (β= −0.2824, se= 0.1995; LRT χ^2^(1)=2.26, p=0.13) (Figure S9). Single-cell RNA sequencing data from the adult human Middle Temporal Gyrus (MTG), as well as brain cell-type transcriptome profiles of adult human (Zhang, et al., 2016) and mouse (Zhang, et al., 2014) brain, showed that *ITIH5* is most highly expressed in the endothelial cells (EC) of the brain (Table S6, Figure S10). This is consistent with its putative role as a brain-blood barrier marker (Daneman, et al., 2010). In the mouse, *Itih5* is one of the 50 most enriched transcripts in brain endothelial cells compared to liver endothelial (3124.8 fold enrichment), or lung endothelial cells (213.3) (Daneman, et al., 2010). *In situ hybridisation* showed that this gene is expressed sporadically and specifically in larger vessels but not in smaller vessels (Daneman, et al., 2010). A more recent characterization of the mouse CNS endothelial cell transcriptome showed *Itih5* is enriched only in adult brain ECs (not in P7 brain ECs, nor in other tissue ECs) (Sabbagh, et al., 2018).

As regards the locus on 2q37, rs7420166 is within the third intron of *BOK* (BCL2-Related ovarian killer) which encodes a member of the BCL2 family, known as essential regulators of the mitochondrial apoptosis pathway (Gross & Katz, 2017; Haschka & Villunger, 2017). The locus also encompasses *BOK-AS1* (BOK antisense RNA 1). In peripheral blood tissue (Westra, et al., 2013; Zhernakova, et al., 2017), rs7420166 and additional SNPs within the associated locus act as eQTLs for *BOK* and 9 other genes in the region (*FARP2, SEPT2, DTYMK, ING5, ATG4B, D2HGDH, AC005104.3, AC114730.7, AC114730.11*) (FDR<0.05). This locus also affects the expression of *BOK* and *DTYMK* in adult cortex (unadjusted p<0.005) (Consortium G., 2015; Ng, et al., 2017).

*BOK* is highly expressed in the central nervous system and multiple brain regions (Table S6) and has been implicated during neuronal apoptosis in mice (D’Orsi, et al., 2016). *BOK* is highly expressed in endothelial cells of the mouse (Table S6), but low expression was detected across human brain cell types (Table S6, Figure S10). In the Allen data from adult post mortem brain, the expression pattern within the PT did not reveal significant hemispheric differences (Figure S9). Developmentally, *BOK* increases in expression levels in the human brain from embryo to mid gestation, and then remains at a steady level into adulthood (Figure S8).

Rs7420166 is located 119kb upstream of *DTYMK*. This gene encodes for an enzyme that catalyzes the phosphorylation of deoxythymidine monophosphate (dTMP) in the deoxythymidine triphosphate (dTTP) synthesis pathway, for DNA synthesis (Huang, et al., 1994). It is most expressed in transformed fibroblasts and lymphocytes, but also present in the brain (most highly in the cerebellum). The expression in the left PT was significantly higher than in the right, in the Allen post mortem brain data (χ^2^(1)=7.18, p= 0.007376) (Figure S9). Developmentally, the expression of *DTYMK* decreases with age, with the highest expression levels at early prenatal time-points (Figure S8).

### 3.4 Testing for broader effects of rs41298373 and rs7420166 on brain asymmetry

Analysis of voxel-wise grey matter volume asymmetry (Figure 2) showed that the effect of rs41298373 was strongest in a cluster (311 voxels, Zmax=7.66, MNI=-40,-34,12) overlapping with the PT and the parietal operculum, as defined in the Harvard-Oxford atlas (Figure 2a). Two additional clusters had effects in the same direction (minor allele A associated with stronger leftward asymmetry): superior temporal gyrus (160 voxels, Zmax=4.49, MNI=-62,-4,4) and temporal pole (65 voxels, Zmax=4.03, MNI=-58,10,-18)(Figure 2). There were also two clusters showing an opposite pattern, i.e. more rightward grey matter volume asymmetry associated with the minor allele of rs41298373, in the parahippocampal gyrus (424 voxels, Zmax=5.45, MNI=-28,-14,-30) and the posterior part of the superior temporal sulcus (112 voxels, Zmax=4.3, MNI=-46,-44,4)(Figure 2). The effect of rs7420166 on grey matter volume asymmetry was largely confined to a single cluster within the PT (Figure 2b, 111 voxels, Zmax=4.22, MNI=-40,-32,14) (minor allele A associated with stronger leftward asymmetry)(Figure 2).

**Figure 2.**
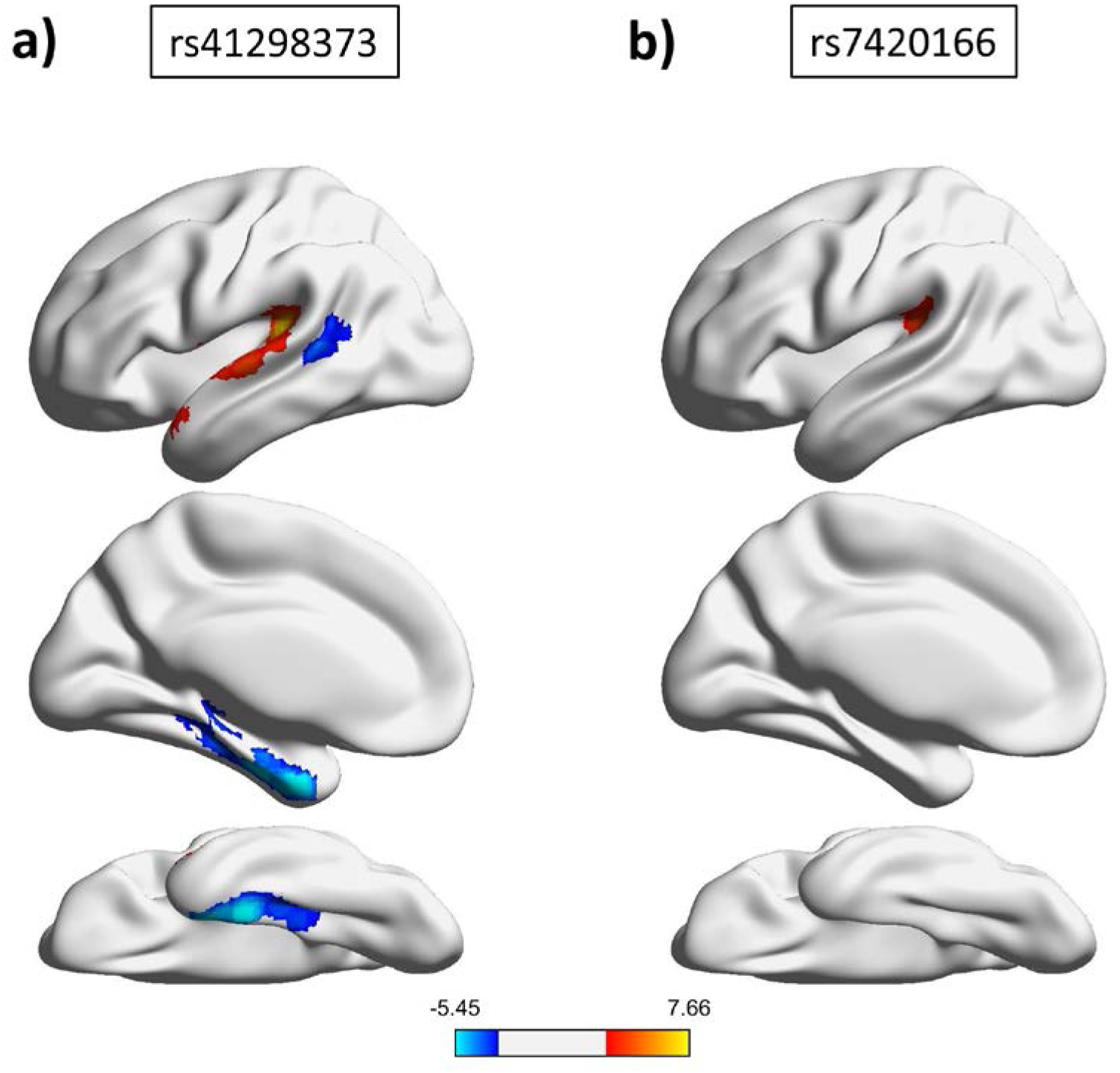
Results of the grey matter VBM AI analysis of **(a)** rs7420166 and **(b)** rs41298373. Z-scores are thresholded at |Z|>3, cluster size >40 voxels, and mapped onto a left-hemisphere surface model for visualization purposes. The direction and strength of the association is indicated by colour (red to yellow = minor allele associated with more leftward asymmetry, blue = minor allele associated with more rightward asymmetry).

In surface-based analysis, there was no significant association of vertex-wise cortical thickness asymmetry with either rs41298373 or rs7420166. Vertex-wise analysis of cortical surface area left-right differences with rs41298373 revealed that the minor allelle A is associated with a more leftward surface area asymmetry in a large cluster of 6380 mm^2^, overlapping with the lower part of the pre-and post-central gyri, the supra-marginal gyrus, the internal part of the superior temporal gyrus covering the PT, Heschl’s gyrus and the posterior part of the insula, and extending to the temporal pole (Figure 3). We also identified a second smaller cluster (2217mm^2^) located in the rostral middle frontal gyrus and showing the opposite pattern (i.e. a more rightward surface area asymmetry associated with the minor allele of rs41298373)(Figure 3). As for rs7420166, we found that the minor allelle is associated with a more leftward cortical surface area asymmetry in a restricted cluster of 2205 mm^2^ overlapping the lower part of the postcentral gyrus, the more anterior part of the supra-marginal gyrus, the PT and the posterior part of Heschl’s gyrus (Figure 3).

**Figure 3.**
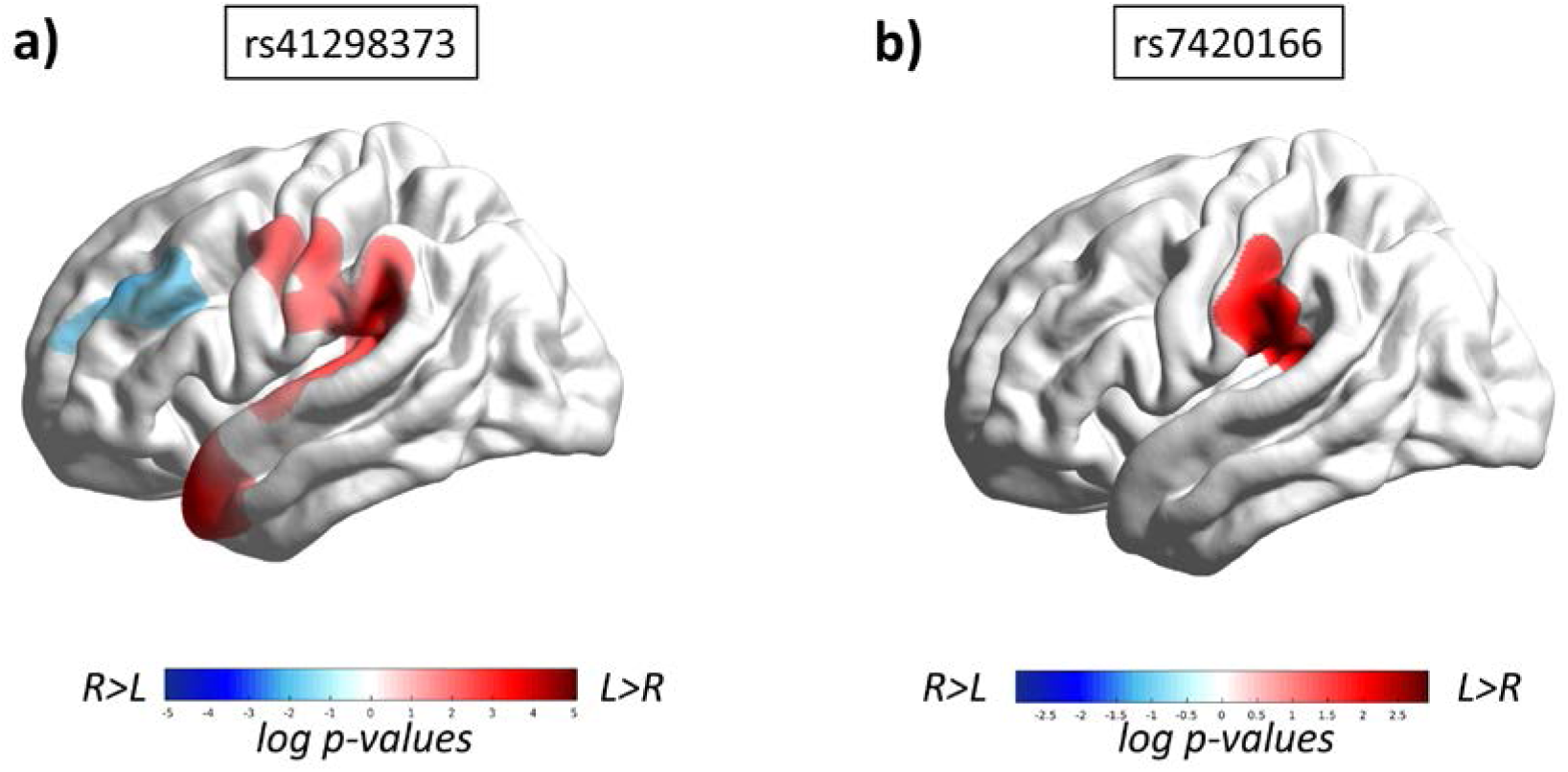
Association analysis of (a) rs41298373 and (b) rs7420166 with vertex-wise cortical surface area left-right differences. Association of minor Allele A with more leftward surface area asymmetry is coded in red, and with more rightward surface area asymmetry in blue. Monte-Carlo simulation corrected log-values p<0.05 are shown.

### 3.5 Gene set enrichment analyses

In gene set enrichment analysis of the GWAS results for PT AI, no gene sets were significantly enriched for association signals after correction for 5,875 tested GO sets.

Since a previous GWAS of PT asymmetry in roughly 3,000 subjects had reported significant enrichment of association in two steroid-related gene sets (Guadalupe, et al., 2015), we also tested these two candidate sets as targeted hypotheses, but found no enrichment of association within ‘steroid hormone receptor activity’ (unadjusted p= 0.68) nor ‘steroid metabolic process’ (unadjusted p= 0.42).

### 3.6 Little evidence for genetic links between PT asymmetry and SCZ, ASD, ADHD, intelligence or educational attainment

Using publicly available GWAS summary statistics from large scale studies of SCZ, ASD, ADHD, intelligence and EA, we found no significant genetic correlations of these traits with PT asymmetry, with a variety of different genetic models and methods (all p>0.05, see Table S8 and Figure S12). Considered separately, the left and right PT volumes were positively genetically correlated with intelligence and EA (Table S8). These genetic correlations were consistent across two out of three methods for calculating them (LDSC and the ldak model of SumHer, Table S8), but were no longer significant after adjusting the left and right PT volumes for TBV, so that these genetic correlations may have largely reflected a general relation of brain size to intelligence and EA. We had adequate power (β>0.80) to detect genetic correlations with PT asymmetry above roughly 0.05-0.15, depending on the trait/model, and as low as 0.025 with the left and right volumes owing to their higher heritabilities than the AI (see Figure S11; Figure S3 Table S3).

In addition, we checked whether either of the two genome-wide significant associations with PT asymmetry which we had detected, i.e. the SNPs rs41298373 and rs7420166, or proxy SNPs for rs7420166, were also associated with any of SCZ, ASD, ADHD, intelligence or EA. There were only two nominally significant associations: between rs41298373 and intelligence (p=0.025) (Savage, et al., 2018), and between three proxy SNPs in high LD with rs7420166 (r^2^>0.80) and EA (all p<0.05, Table S9). Note that not all of the various GWAS summary statistics included data on these SNPs or their proxies for the purposes of this look-up (see Table S9).

## 4 Discussion

It has been a longstanding hypothesis that asymmetry of the PT is linked genetically to variation in human cognition and disorder susceptibility (Sommer, Ramsey, Kahn, Aleman, & Bouma, 2001; Crow, 1993), given evidence for associations at the phenotypic level (see Introduction). Here we leveraged molecular genetic and neuroimaging data, which have only become available on a sufficiently large scale very recently, to investigate this question from a polygenic perspective. Overall, our analysis yielded novel insights into the genetic contributions to PT asymmetry, but did not support a clear role of these in cognitive variation and disorder susceptibility, at least that could be detected in current sample sizes.

### Genetics of PT asymmetry

Within the UK Biobank dataset, we showed that asymmetry of the PT has a modest but significant SNP-based heritability, estimated at 14%. In principal, a heritable component to PT asymmetry may arise because some genetic influences are quantitatively different on the left and right volumes of this structure, and we found evidence to support this, as the genetic correlation between left and right PT volumes was significantly less than 1. However, PT asymmetry may itself be considered a primary brain phenotype, not merely arising from different effects on left and right. This conception would be consistent with the early appearance of PT asymmetry in development *in utero*, and its association with disorders when disrupted. Thus PT asymmetry may be analogous to the placement of the visceral organs on the left-right body axis (heart, lungs etc.), which arises due to a well-regulated genetic-developmental program in most people, but can be disrupted by various genetic mutations and environmental insults (Deng, Xia, & Deng, 2015). In other words, there are probably some genetic influences on primary development of the brain’s left-right axis, as well as other genetic influences that can have different effects on the left and right sides, which may come into play after the two sides have already been sent on different developmental trajectories. Asymmetry is also likely to be affected and maintained by interactions between the two hemispheres throughout development and adulthood. Studies in which gene expression has been contrasted between the left and right sides of the embryonic and foetal central nervous systems have provided support for a genetic developmental program underlying brain asymmetry (Sun, et al., 2005; Ocklenburg, et al., 2017; de Kovel C. G., et al., 2017; de Kovel C. G., Lisgo, Fisher, & Francks, 2018), although they could not specifically target the primitive PT itself.

A nonsynonymous variant within *ITIH5* was significantly associated with PT asymmetry, for which the minor allele increased the left PT volume, but did not affect the right. VBM association analysis indicated that this variant affects asymmetry of the PT and the directly neighbouring parietal opercular cortex most strongly, but also extends to other temporal regions. Interestingly, the effect on asymmetry is in opposite directions in the PT and nearby middle temporal gyrus, suggesting that this variant affects the relative distribution of cortical tissue between these two left-hemisphere regions, and/or inter-hemispheric differences in the sylvian fissure’s position and ending. In vertex-wise, surface-based analysis using hemispheric co-registration, the effect is strongest in the PT area but also extends along the superior temporal surface area.

The non-synonymous variant in *ITIH5* occurs at an evolutionarily conserved site, and the minor allele, which is the derived rather than ancestral allele, results in an amino acid change that is predicted to affect protein function and to be deleterious (i.e. selected against). Functional experiments will be needed to confirm whether the protein is impacted by this variant and how this might affect regional cortical development. From the perspective of language evolution, it is intriguing to note that a variant in the modern human population which leads to a larger left PT volume might also be selected against. However, any deleteriousness arising from this allele is not necessarily mediated through its effects on left PT volume or asymmetry. *ITIH5* is also expressed in many other non-brain tissues, with an apparent role in stabilizing the extracellular matrix.

From various data sources we found that *ITIH5* is expressed at moderate levels in the brain, in a mostly constant manner across regions and developmental stages, and with high expression in endothelial cells, consistent with a potential role as a blood-brain-barrier protein. There was a non-significant trend toward lower right-than left-side expression in the adult PT, in the Allen brain data, but the number of samples was too low to yield a conclusive result regarding expression levels of this gene. Overall, the available expression data for ITIH5 do not point to a clear explanation for the regional specificity of this gene-brain association, which may arise due to interactions of ITIH5 with other hemisphere-and region-specific factors.

Another insight into the genetics of PT asymmetry involved a locus on 2q37 which affects the expression level of *BOK*, a gene which has a role in neuronal apoptosis (D’Orsi, et al., 2016), and also the kinase *DTYMK* at the same locus, which is involved in DNA synthesis and is expressed at a higher level on the left PT than the right, in post mortem adult tissue. The VBM association analysis indicated that this effect is specific to asymmetry of the PT region. However, vertex-wise analysis indicated an association with cortical surface area asymmetry that is somewhat more diffuse, covering a broader region of the internal part of the superior temporal gyrus, including the PT. This may be due to more accurate inter-hemispheric alignment in the surface-based analysis as compared to the voxel-based one. Again, interactions of this gene with other hemisphere and region-specific factors may explain this restricted gene-brain association pattern.

### Relation to SCZ, ASD, ADHD, intelligence, EA

We had 80% statistical power to detect genetic correlations with PT asymmetry larger than roughly 0.05-0.10 for SCZ, intelligence and EA, or 0.15 for ASD and ADHD, given the sample sizes used to derive the GWAS summary statistics for these traits, and their SNP-based heritabilities. Despite this, no significant genetic correlations were detected between PT asymmetry and these traits or disorders. One possibility is that previous studies which have shown phenotypic associations between PT asymmetry and SCZ and ASD in particular, have reported false positive findings based on limited sample sizes and/or publication bias in meta-analysis. In general there is an urgent need for the field to increase sample sizes in brain-disorder association studies. As regards ADHD, intelligence and EA, we are not aware of studies which have reported direct phenotypic correlations with PT asymmetry, but such reports have been made for dyslexia, while reading ability shares genetic contributions with ADHD, intelligence and EA (see Introduction). We await GWAS studies of reading measures in more than 10,000 subjects, to perform future genetic correlation analyses with PT asymmetry.

It is also possible that phenotypic associations between PT asymmetry and cognitive/psychiatric disorders may not be genetically driven, but caused by environmental factors. In line with this, a previous study also found no genetic overlap between SCZ and eight brain volume measures (mainly subcortical, and not with reference to asymmetry), despite robust phenotypic associations having been reported of these volumes with SCZ (Franke, et al., 2016).

A third possibility is that genetic correlations do exist between PT asymmetry and some of SCZ, ASD, ADHD, intelligence, EA, but that the correlations are too low to have been detected with the present sample sizes. The GWAS for PT asymmetry in the present study was based on over 18,000 subjects, while the disorder GWAS were based on over 20,000 cases each, the intelligence GWAS over 250,000 individuals, and the EA GWAS over 750,000 individuals. As sample sizes increase further in the coming years, it may still be possible to detect genetic correlations of PT asymmetry with some of these traits, of the order of 0.05 or lower. Regardless, we can already conclude that PT asymmetry probably does not have a substantial genetic correlation with any of these traits.

We also explored the genetic correlations of the left and right PT volumes separately with SCZ, ASD, ADHD, intelligence, EA. In this case, as the left and right volume measures had higher heritabilities than the AI, we had 80% power to detect genetic correlations as low as 0.05. We found that intelligence and EA had positive genetic correlations with both left and right PT volumes to similar extents. However, after adjusting the left and right PT volumes for TBV, the genetic correlations with intelligence and EA were largely diminished and non-significant. A previous study reported that average total cortical surface area is genetically correlated with EA (rg=0.213, p=9.47×10^−13^) and general cognitive function (rg=0.2221, p=3.94×10^−8^) (Grasby, et al., 2018). Hence, the genetic correlations of left and right PT volumes with intelligence and EA, which we detected in the present study, are likely to reflect global effects of brain size, rather than region-specific effects, and are not linked to laterality.

Regarding the two individual loci which were significantly associated with PT asymmetry in the GWAS for this trait, we did not find evidence that they were associated with SCZ, ASD, or ADHD. The non-synonymous SNP in *ITIH5* was nominally associated with intelligence (unadjusted P=0.02) with a very small effect size (standard beta= 0.011). Thus, it is possible that this locus does impact intelligence, although a P value of 0.02 based on a study of 269,867 individuals means that this could only be clarified in a much larger, future dataset. Rs7420166 was not available in the intelligence and EA GWASes, but three SNPs in high LD with it (r^2^>0.8: rs114606940, rs73123528, rs77278412) were nominally associated with EA (unadjusted Ps <0.05). Again, these effects (beta=0.004) were very small, detectable only in a dataset of 766,345 individuals.

### Limitations

Limitations with regard to sample size, statistical power, and the lack of large scale GWAS for dyslexia have already been discussed above. As regards dyslexia, the phenotypic association with PT asymmetry has been reported to be limited to boys (see Introduction), which means that further sex-limited genetic correlation analysis may be warranted. For the present study, the brain imaging data in the UK Biobank would be reduced to groups of roughly 8,500-9,000 if split by sex, which is too low to perform well powered genetic correlation analysis with other datasets by use of summary statistics, for a trait such as PT asymmetry with only 14% heritability. Also, the GWAS summary statistics for SCZ, ASD, intelligence and EA were not available split by sex. Although sex-stratified GWAS summary statistics were available for ADHD, the power to detect genetic correlation of PT asymmetry with this disorder would not have been adequate in sex-split analysis. In this study we controlled for sex as a covariate, and also found no evidence for a sex-specific effect of the top loci associated with PT asymmetry.

The Harvard-Oxford atlas, which we used to define the PT, was derived from manual segmentations of sets of reference brain images (Destrieux, Fischl, Dale, & Halgren, 2010; Goldstein, et al., 1999; Goldstein, et al., 2007). It therefore contained asymmetrical definitions for regions that showed asymmetry in the reference dataset (including the PT). Accordingly, the measurement of average PT asymmetry in the UK Biobank dataset would partly reflect left-right differences present in the atlas. Nonetheless, the use of a ‘real-world’ asymmetrical atlas, rather than an artificially created symmetrical atlas, was appropriate for our primary purpose which was the GWAS of PT asymmetry in as large a sample as possible. This is because regional identification was likely to be more accurate for structures that were asymmetrical both in the atlas and, on average, in the UK Biobank dataset, and the GWAS analysis is based on comparing relative levels of asymmetry between genotype groups, rather than measuring absolute asymmetry levels. In addition, we followed up the most significant SNP associations with PT asymmetry that arose in the GWAS, by performing brain-wide grey matter asymmetry VBM association analysis, this time based on a symmetrized template and no regional atlas definitions, which produced results that were highly consistent with the GWAS analysis. We further investigated vertex-wise effects without regard to any regional atlas definitions, and using a co-registration approach to improve interhemispheric correspondence.

### Future perspectives

This study provided various novel insights into the genetic contributions to PT asymmetry, but did not find evidence that these are substantially shared with cognitive variation or susceptibility to psychiatric disorders. Future studies based on even larger samples may be able to tease out very subtle genetic correlations.

Comparative studies have shown that chimpanzees have a leftward asymmetry of the Sylvian fissure and PT, while several old world monkey species do not (Yeni-Komshian & Benson, 1976; Lyn, et al., 2011). A recent study reported the first population-level leftward asymmetry of the PT in a non-hominin primate (olive baboon monkeys), dating the origin of this cerebral specialization as early as 30-40 million years ago (Marie, et al., 2018). Therefore PT asymmetry is not a uniquely human feature. The involvement of the PT in language processing, which usually shows left-side dominance, probably arose through adaption of an already-lateralized brain infrastructure. Nonetheless, future studies may interrogate the genetics of PT asymmetry from the perspective of genomic conservation and signals of selection.

## Acknowledgements

This research was conducted using the UK Biobank Resource under Application Number 16066, with Clyde Francks as the principal applicant. AC-C was funded by a grant to CF from the Netherlands Organization for Scientific Research (NWO)(054-15-101), and B.M, N.T.M., A.P., and F.C by a grant from the French National Research Agency (ANR, grant No. 15-HBPR-0001-03), as part of the FLAG-ERA consortium project ‘MULTI-LATERAL’, a Partner Project to the European Union’s Flagship Human Brain Project. Additional support was from the Max Planck Society (Germany).

