## Supplementary material for "Genetic effects on *planum temporale* asymmetry and their limited relevance to neurodevelopmental disorders, intelligence or educational attainment"

### Supplementary tables

**Table S1: Overview of covariates included in the regression to prepare residualized PT and TBV measures. fID= UK Biobank variable ID.**

| Covariate | fID | Transformation | Counts | Range (Min; Max) | Mean | SD |
|---|---|---|---|---|---|---|
| age | - | (date imaging) - (year of birth+month of birth)<br>date imaging fID = f.53.2.0<br>year of birth fID = f.34.0.0<br>month of birth fID = f.52.0.0 | | 45.2; 80.7 | 63.33 | 7.45 |
| zage2 | - | $(age-mean(age))^2$ | | 0.0007; 328.6 | 55.47 | 58.66 |
| sex | f.22001.0.0 | - | Males = 8631<br>Females= 9426 | | | |
| PC1 | f.22009.0.1 | - | | -18.05; -6.26 | -12.40 | 1.63 |
| PC2 | f.22009.0.2 | - | | -1.89; 9.35 | 3.78 | 1.51 |
| PC3 | f.22009.0.3 | - | | -7.68; 4.40 | -1.64 | 1.59 |
| PC4 | f.22009.0.4 | - | | -9.51; 12.68 | 1.37 | 2.85 |
| PC5 | f.22009.0.5 | - | | -14.8; 24.66 | -1.26 | 6.47 |
| PC6 | f.22009.0.6 | - | | -6.53; 5.93 | -0.46 | 1.66 |
| PC7 | f.22009.0.7 | - | | -12.05; 8.02 | 0.21 | 1.85 |
| PC8 | f.22009.0.8 | - | | -8.28; 15.24 | -0.67 | 1.99 |
| PC9 | f.22009.0.9 | - | | -33.19; 11.91 | 0.72 | 3.92 |
| PC10 | f.22009.0.10 | - | | -7.68; 9.58 | 0.53 | 2.30 |
| array | - | - | UKBB = 16274<br>UKBL = 1783 | | | |
| Scanner lateral X brain position | f.25756.2.0 | | | -9.85; 17.28 | 0.43 | 2.57 |
| Scanner transverse Y brain position | f.25757.2.0 | - | | 53; 96 | 63.59 | 5.49 |
| Scanner longitudinal Z brain position | f.25758.2.0 | - | | -113.3; 124.5 | -17.92 | 31.84 |

**Table S2: Summary statistics for PT measures and TBV.** N indicates the number of subjects after residualizing for covariates (see main manuscript).

| Phenotype | Range (Min; Max) | Mean | SD | N |
|---|---|---|---|---|
| AI | -0.4971; 0.9852 | 0.236 | 0.190 | 18049 |
| L | 869; 3985 | 2056 | 479.835 | 18037 |
| R | 664.7; 2881 | 1607 | 316.062 | 18039 |
| TBV | 828168; 1612260 | 1167877 | 111175.100 | 18053 |

**Table S3: Heritability estimates for PT measures and TBV, using GREML.**

| Phenotype | $h^2$ | se | P | N |
|---|---|---|---|---|
| AI | 0.139 | 0.035 | $1.38\times10^{-5}$ | 17205 |
| L | 0.454 | 0.037 | 0 | 17185 |
| R | 0.414 | 0.036 | 0 | 17194 |
| TBV | 0.717 | 0.036 | 0 | 17209 |

**Table S4: Phenotypic and genetic correlations between PT measures and TBV, after residualizing for covariate effects.** The covariates are shown in Table S1. Genetic correlations were calculated using bivariate GREML.*The genetic correlation ρ was tested against the null hypothesis that ρ=1, otherwise the null hypothesis was that ρ=0.

| | | Phenotypic correlation | | Genetic correlation | | |
|---|---|---|---|---|---|---|
| | | r | P | ρ | se | P |
| L | R | 0.590 | 0 | 0.858 | 0.061 | 0.00889 (*) |
| L | TBV | 0.527 | 0 | 0.523 | 0.039 | 0 |
| R | TBV | 0.557 | 0 | 0.541 | 0.038 | 0 |
| AI | TBV | 0.071 | 9.40E-22 | 0.133 | 0.083 | 0.053 |

**Table S5: SNPs associated at a genome-wide significant level with either the PT AI, left PT, or right PT.** The $r^2$ indicates linkage disequilibrium with the lead SNP, as calculated within the UKB imaging dataset. A1: effect allele, the direction of the T value indicates the effect of having each extra A1 allele. Minor: minor allele. MAF: minor allele frequency. Location: chromosome and position in the genome (hg19). P values in **bold font** are significant in the genome-wide context.

| Lead SNP GWAS | r2 | SNP identity | A0 | A1 | Minor | MAF | Location | AI | | Left | | Right | |
|---|---|---|---|---|---|---|---|---|---|---|---|---|---|
| | | | | | | | | T | P | T | P | T | P |
| rs7420166 | 0.743 | rs7576339 | G | A | A | 0.248 | 2:242490882 | 5.6493 | **$1.64\times10^{-8}$** | 2.7173 | 0.006589 | -2.6578 | 0.007872 |
| | 0.877 | rs77278412 | T | A | A | 0.238 | 2:242492109 | 5.482 | **$4.26\times10^{-8}$** | 3.0294 | 0.002454 | -2.0975 | 0.035967 |
| | 0.886 | rs114606940 | C | T | T | 0.239 | 2:242496904 | 5.5256 | **$3.33\times10^{-8}$** | 3.1175 | 0.001827 | -2.0855 | 0.037034 |
| | 0.869 | rs115841724 | C | T | T | 0.236 | 2:242502394 | 5.4984 | **$3.89\times10^{-8}$** | 3.1999 | 0.001377 | -1.8901 | 0.058762 |
| | 0.898 | rs146946923 | C | CTGTA | CTGTA | 0.237 | 2:242505574 | 5.5949 | **$2.24\times10^{-8}$** | 3.2251 | 0.001262 | -1.8936 | 0.058291 |
| | - | **rs7420166** | G | A | A | 0.259 | 2:242506733 | 6.1577 | **$7.54\times10^{-10}$** | 3.7038 | 0.000213 | -1.9504 | 0.051145 |
| rs41298373 | - | **rs41298373** | G | A | A | 0.101 | 10:7622009 | 7.9478 | **$2.01\times10^{-15}$** | 6.1817 | **$6.48\times10^{-10}$** | -1.4971 | 0.134385 |
| rs12932673 | - | **rs12932673** | C | G | G | 0.189 | 16:86709731 | 1.9413 | 0.05224 | -2.757 | 0.00584 | -5.5896 | **$2.31\times10^{-8}$** |
| | 0.979 | rs10048146 | A | G | G | 0.194 | 16:86710660 | 1.9708 | 0.048764 | -2.7025 | 0.006888 | -5.5769 | **$2.48\times10^{-8}$** |

**Table S6: Summary information for genes of interest which arose from the PT AI GWAS in the UK Biobank dataset.**

| Locus | Gene symbol | Gene name | GTEx | | BRAINEAC | Allen Brain Atlas (Expression per region, averaged across probes and samples) | | Allen Brain Atlas: single-cell RNAseq | BrainRNASeq | |
|---|---|---|---|---|---|---|---|---|---|---|
| | | | Highest tissue (TPM) | Highest brain (TPM) | Expression range (log2) | Highest expression | Lowest expression | Highest expression cell cluster (group median CPM) | Human cell type with highest expression (FPKM) | Mouse cell type with highest expression (FPKM) |
| 10p14 | *ITIH5* | Inter-alpha-trypsin inhibitor, heavy chain 5 | Adipose subcutaneous (105.5) | Putamen, basal ganglia (9.96) | Medulla (6) Ceberellum (5) | Gracile nucleus left (9.34) | Dentate gyrus left (3.76) | Endo L2-6 NOSTRIN (546.2) | Endothelial (18.3) | Oligodendrocite precurso cells (15.3) |
| 2q37.3 | *BOK* | BCL2-Related ovarian killer | Brain spinal cord, cervical c-1 (175) | Spinal cord, cervical c-1 (175) | White matter (8.8) Cerebellum (8.8) | Red nucleus left (11.27) | Dentate gyrus left (2.99) | 0 | Oligodendro cites (0.81) | Endothelial (94.94) |
| 2q37.3 | *DTYMK* | Deoxythymidylate Kinase | Cells - EBV-transformed lymphocytes: (50.16) | Brain - Cerebellar Hemisphere (22.37) | - | Central Glial Substance (10.08) | CA2 Field Right (5.587) | Exc L5-6 THEMIS FGF10 (0.072) | Fetal Astrocytes (10.27) | Oligodendrocite precursor cells (41.69) |

**Table S7: GWAS studies of psychiatric, cognitive and behavioural traits of interest, for which summary statistics are publicly available and were used to calculate genetic correlations with the PT measures.** Prev=prevalence of disorder in the general population. For each trait, the sample size and prevalence were extracted from the original reference. The $h^2_{SNP}$(SE) estimates were calculated using LDSC and SumHer (gcta and ldak models). These parameters were used in the power calculations.

| PMID | Reference | Trait | N | Ncases | Ncntr | Prev | $h^2_{SNP}$(SE) | | |
|---|---|---|---|---|---|---|---|---|---|
| | | | | | | | LDSC | SumHer -gcta | SumHer -ldak |
| 25056061 | Ripke et al. (2014) (44) | SCZ | | 36,989 | 113,075 | 0.01 | 0.18 (0.009) | 0.35 (0.014) | 0.43 (0.015) |
| - | Demontis et al. (2017) (46) | ADHD | | 18,382 | 27,969 | 0.05 | 0.21 (0.015) | 0.22 (0.010) | 0.32 (0.014) |
| - | Grove et al. (2017) (45) | ASD | | 19,099 | 34,194 | 0.012 | 0.12 (0.017) | 0.11 (0.010) | 0.17 (0.014) |
| 30038396 | Lee et al. (2018) (49) | EA | 766,345 | | | - | 0.11 (0.003) | 0.14 (0.003) | 0.17 (0.004) |
| 29942086 | Savage et al. (2018) (48) | Intelligence | 269,867 | | | - | 0.18 (0.008) | 0.26 (0.007) | 0.35 (0.008) |

**Table S8: Genetic correlations (ρ) between PT volume phenotypes and other traits of interest**

| PT | Psychiatric disorder/ Cognitive trait | LDSC | | SumHer-gcta | | SumHer-ldak | |
|---|---|---|---|---|---|---|---|
| | | ρ (SE) | P | ρ (SE) | P | ρ (SE) | P |
| AI | Intelligence | 0.02 (0.08) | 0.81 | 0.036 (0.09) | 0.67 | 0.05 (0.12) | 0.69 |
| AI | EA | -0.038 (0.06) | 0.51 | -0.049 (0.09) | 0.57 | -0.18 (0.14) | 0.2 |
| AI | ASD | 0.206 (0.15) | 0.17 | 0.047 (0.03) | 0.13 | 0.026 (0.09) | 0.76 |
| AI | ADHD | 0.049 (0.12) | 0.67 | 0.055 (0.03) | 0.08 | 0.095 (0.09) | 0.31 |
| AI | SCZ | -0.013 (0.08) | 0.86 | 0.026 (0.02) | 0.24 | 0.07 (0.07) | 0.29 |
| L | Intelligence | 0.154 (0.05) | $3.40\times10^{-3}$ | 0.112 (0.06) | 0.08 | 0.154 (0.05) | $1.40\times10^{-3}$ |
| L | EA | 0.119 (0.04) | $1.50\times10^{-3}$ | 0.096 (0.06) | 0.12 | 0.101 (0.05) | 0.03 |
| L | ASD | 0.119 (0.08) | 0.16 | 0.015 (0.02) | 0.49 | -0.016 (0.04) | 0.68 |
| L | ADHD | -0.092 (0.06) | 0.16 | 0.013 (0.02) | 0.46 | 0.002 (0.03) | 0.95 |
| L | SCZ | -0.016 (0.04) | 0.71 | 0.019 (0.02) | 0.21 | 0.007 (0.03) | 0.79 |
| R | Intelligence | 0.16 (0.05) | $2.20\times10^{-3}$ | 0.097 (0.06) | 0.13 | 0.149 (0.05) | $2.60\times10^{-3}$ |
| R | EA | 0.157 (0.04) | $9.10\times10^{-6}$ | 0.128 (0.06) | 0.03 | 0.182 (0.05) | $7.30\times10^{-5}$ |
| R | ASD | 0.03 (0.07) | 0.68 | -0.007 (0.02) | 0.71 | -0.025 (0.04) | 0.49 |
| R | ADHD | -0.148 (0.06) | 0.02 | -0.024 (0.02) | 0.16 | -0.031 (0.03) | 0.33 |
| R | SCZ | -0.018 (0.05) | 0.7 | 0.005 (0.01) | 0.73 | -0.016 (0.03) | 0.53 |

**Table S9: Lookup of lead SNPs which were significantly associated with PT asymmetry in the GWAS, in publicly available GWAS results for other traits of interest.** The $r^2$ indicates linkage disequilibrium with the lead SNP, in case proxy SNPs were needed to make the lookup, as calculated within the UKB imaging dataset. Pos: chromosomic position of the SNP in the hg19 human reference genome. INFO: imputation quality score. OR/Beta/stdBeta: association statistic.

| Lead SNP from PT AI GWAS | $r^2$ | Proxy SNP | Chr | Pos | Trait | P | INFO | min INFO | OR | Beta | stdBeta | SE |
|---|---|---|---|---|---|---|---|---|---|---|---|---|
| rs7420166 | 0.89 | rs114606940 | 2 | 242496904 | Intelligence | 0.458 | | 0.81 | | | -0.002 | 0.003 |
| | | | | | EA | 0.034 | | | | 0.004 | | 0.002 |
| | | | | | ASD | 0.609 | 0.92 | | 1.01 | | | 0.017 |
| | | | | | ADHD | 0.871 | 0.92 | | 1.00 | | | 0.019 |
| | | | | | SCZ | 0.060 | 0.81 | | 0.97 | | | 0.014 |
| | 0.87 | rs115841724 | 2 | 242502394 | Intelligence | 0.567 | | 0.78 | | | -0.002 | 0.003 |
| | | | | | EA | 0.093 | | | | 0.003 | | 0.002 |
| | | | | | ASD | 0.827 | 0.88 | | 1.00 | | | 0.017 |
| | | | | | ADHD | 0.361 | 0.88 | | 0.98 | | | 0.019 |
| | | | | | SCZ | 0.083 | 0.78 | | 0.98 | | | 0.014 |
| | 0.90 | rs146946923 | 2 | 242505574 | Intelligence | - | | | | | | |
| | | | | | EA | - | | | | | | |
| | | | | | ASD | 0.764 | 0.89 | | 1.01 | | | 0.017 |
| | | | | | ADHD | 0.420 | 0.89 | | 0.98 | | | 0.019 |
| | | | | | SCZ | - | | | | | | |
| | 0.87 | rs73123528 | 2 | 242504144 | Intelligence | 0.268 | | 0.75 | | | 0.003 | 0.003 |
| | | | | | EA | 0.045 | | | | 0.004 | | 0.002 |
| | | | | | ASD | 0.789 | 0.88 | | 1.00 | | | 0.017 |
| | | | | | ADHD | 0.440 | 0.88 | | 1.01 | | | 0.019 |
| | | | | | SCZ | 0.158 | 0.79 | | 1.02 | | | 0.013 |
| | 1 | rs7420166 | 2 | 242506733 | Intelligence | - | | | | | | |
| | | | | | EA | - | | | | | | |

| | | | | | | | | | | | | |
|---|---|---|---|---|---|---|---|---|---|---|---|---|
| | | | | | ASD | 0.601 | 0.88 | | 1.01 | | | 0.017 |
| | | | | | ADHD | 0.762 | 0.88 | | 0.99 | | | 0.019 |
| | | | | | SCZ | 0.190 | 0.77 | | 0.98 | | | 0.014 |
| | 0.86 | rs74562536 | 2 | 242500151 | Intelligence | 0.648 | | 0.80 | | | -0.002 | 0.003 |
| | | | | | EA | 0.061 | | | | -0.004 | | 0.002 |
| | | | | | ASD | 0.561 | 0.90 | | 1.01 | | | 0.017 |
| | | | | | ADHD | 0.631 | 0.91 | | 0.99 | | | 0.019 |
| | | | | | SCZ | 0.062 | 0.81 | | 0.97 | | | 0.014 |
| | 0.88 | rs76812504 | 2 | 242503739 | Intelligence | 0.207 | | 0.77 | | | -0.004 | 0.003 |
| | | | | | EA | 0.052 | | | | -0.004 | | 0.002 |
| | | | | | ASD | 0.751 | 0.89 | | 1.01 | | | 0.017 |
| | | | | | ADHD | 0.486 | 0.89 | | 0.99 | | | 0.019 |
| | | | | | SCZ | 0.056 | 0.81 | | 0.97 | | | 0.014 |
| | 0.88 | rs77278412 | 2 | 242492109 | Intelligence | 0.849 | | 0.82 | | | -0.001 | 0.003 |
| | | | | | EA | 0.033 | | | | -0.004 | | 0.002 |
| | | | | | ASD | 0.646 | 0.92 | | 1.01 | | | 0.017 |
| | | | | | ADHD | 0.894 | 0.93 | | 1.00 | | | 0.019 |
| | | | | | SCZ | 0.073 | 0.82 | | 0.98 | | | 0.014 |
| rs41298373 | - | rs41298373 | 10 | 7622009 | Intelligence | 0.025 | | 0.62 | | | 0.011 | 0.005 |
| | | | | | EA | 0.201 | | | | 0.004 | | 0.003 |
| | | | | | ASD | 0.765 | 0.94 | | 1.01 | | | 0.023 |
| | | | | | ADHD | 0.693 | 1.00 | | 0.99 | | | 0.025 |
| | | | | | SCZ | - | | | | | | |

#### Supplementary figures

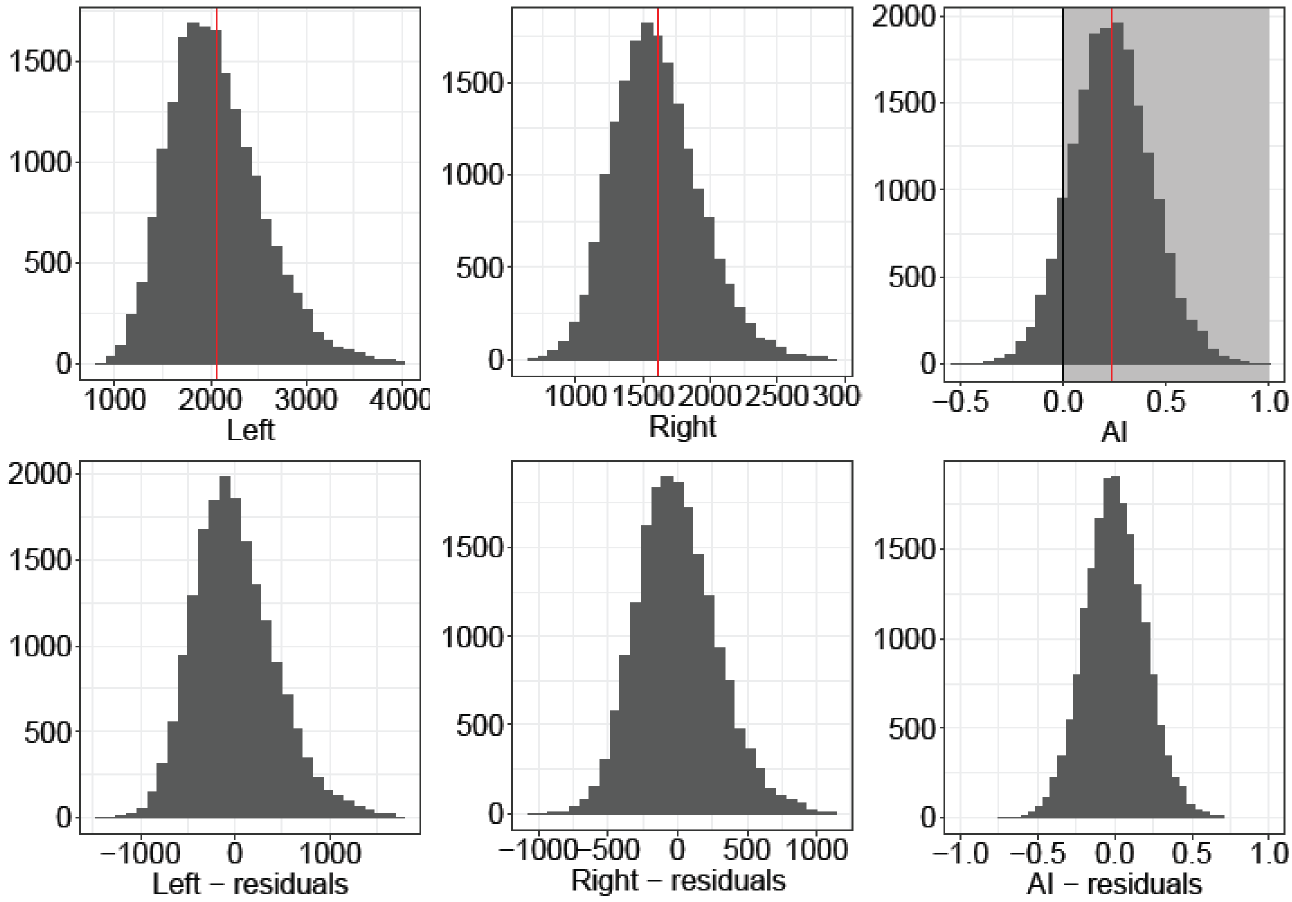


**Figure S1: Distributions of the planum temporale (PT) volume phenotypes before (top) and after (bottom) accounting for covariates.** The red solid lines indicate the mean of each phenotype. The black solid line indicates the point at which AI=0, and the shaded area highlights that the PT is leftward asymmetrical for most people.

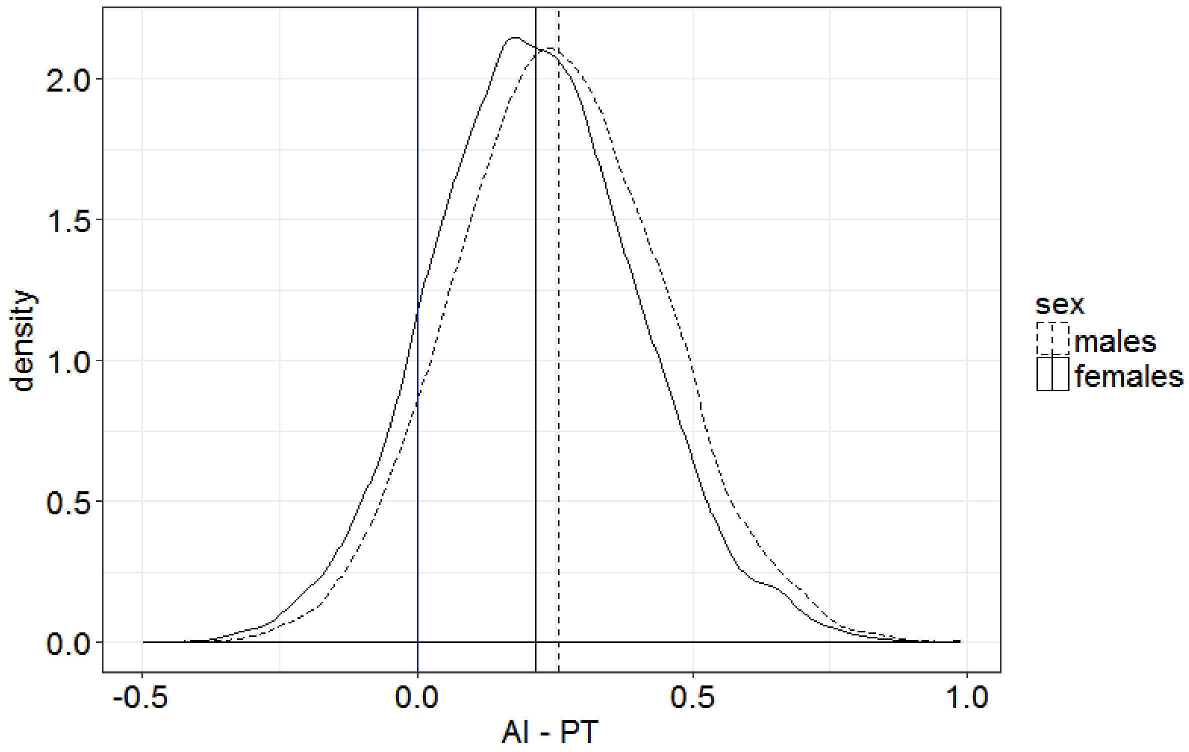


**Figure S2: Density plot of the planum temporale asymmetry index (PT AI) separately by sex.** The blue line indicates the symmetrical AI. The solid lines indicate the mean per group (solid=females, dotted=males).

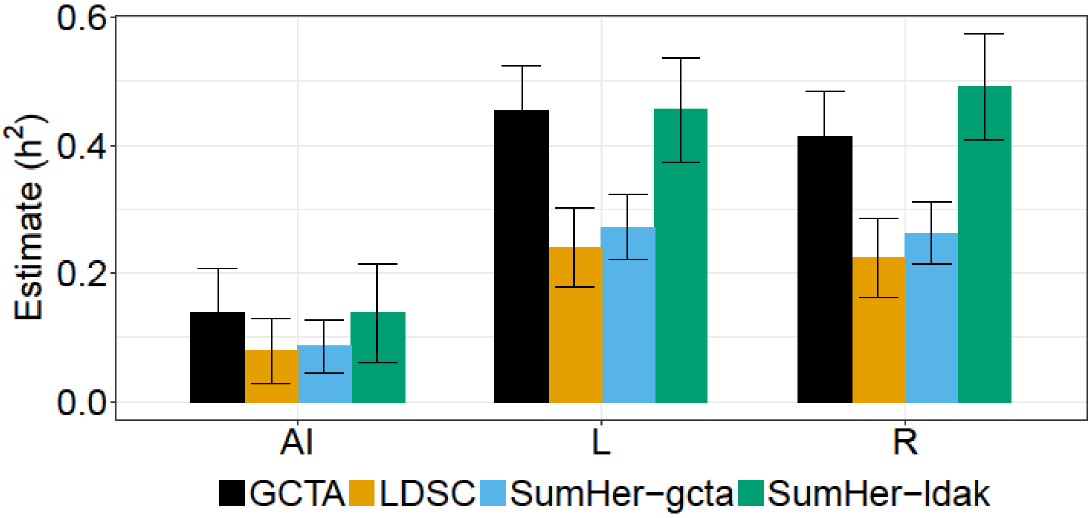


**Figure S3: SNP-based heritability estimates for PT volume measures as calculated with different methods.** The lines indicate the 95 % confidence intervals for the estimates.

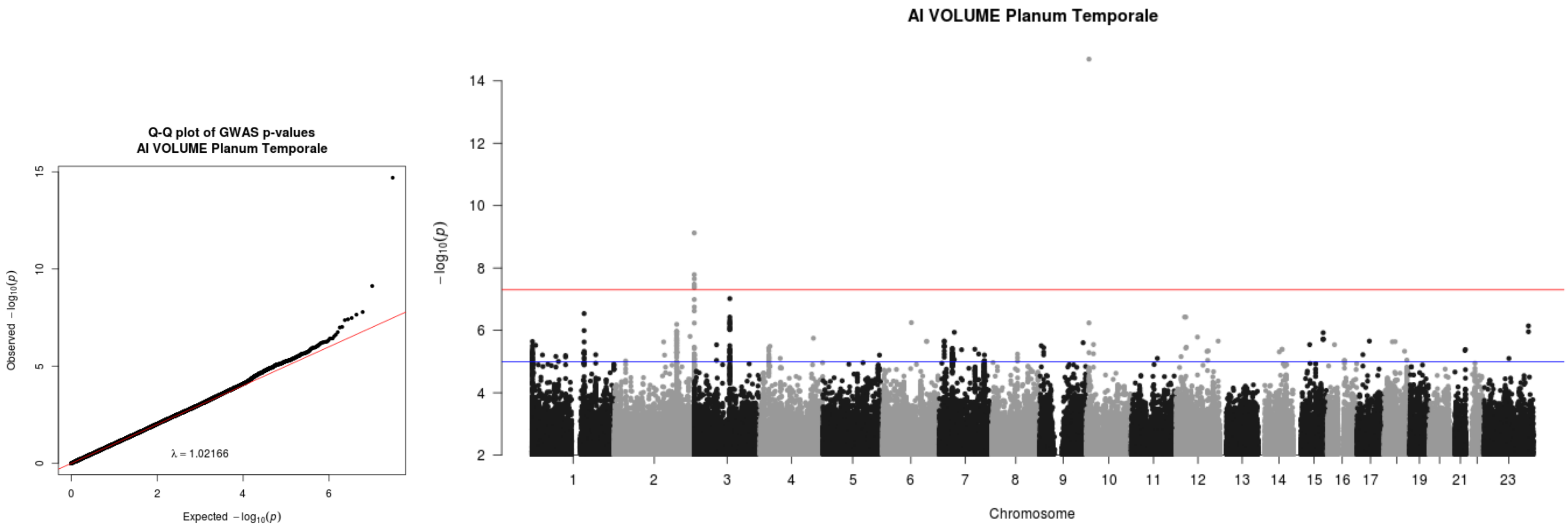


**Figure S4: QQ plot and Manhattan plot from the GWAS for PT AI.**

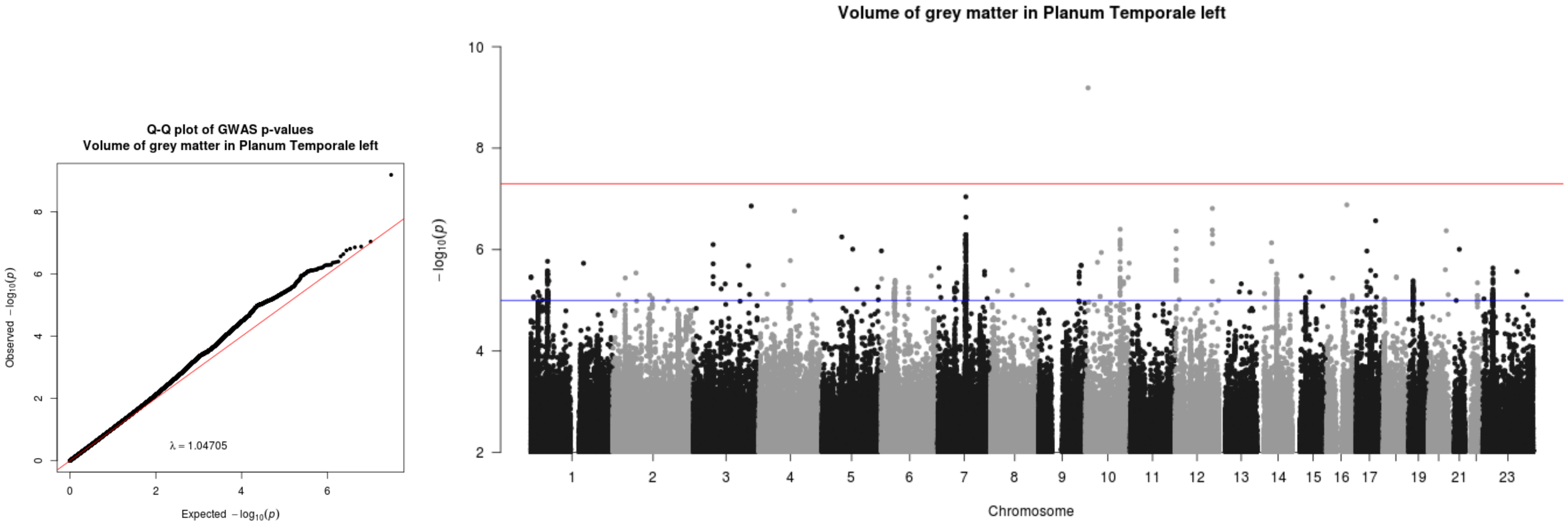


**Figure S5: QQ plot and Manhattan plot from the GWAS for the left PT.**

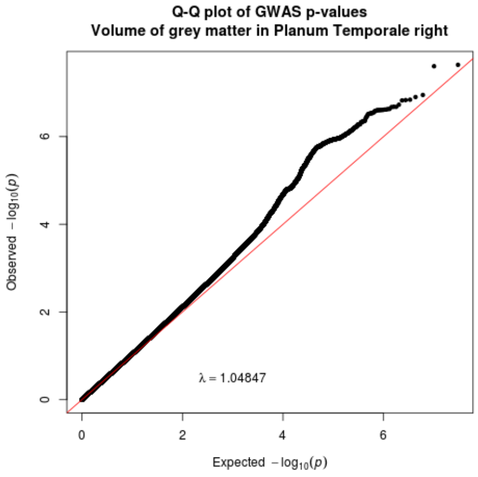


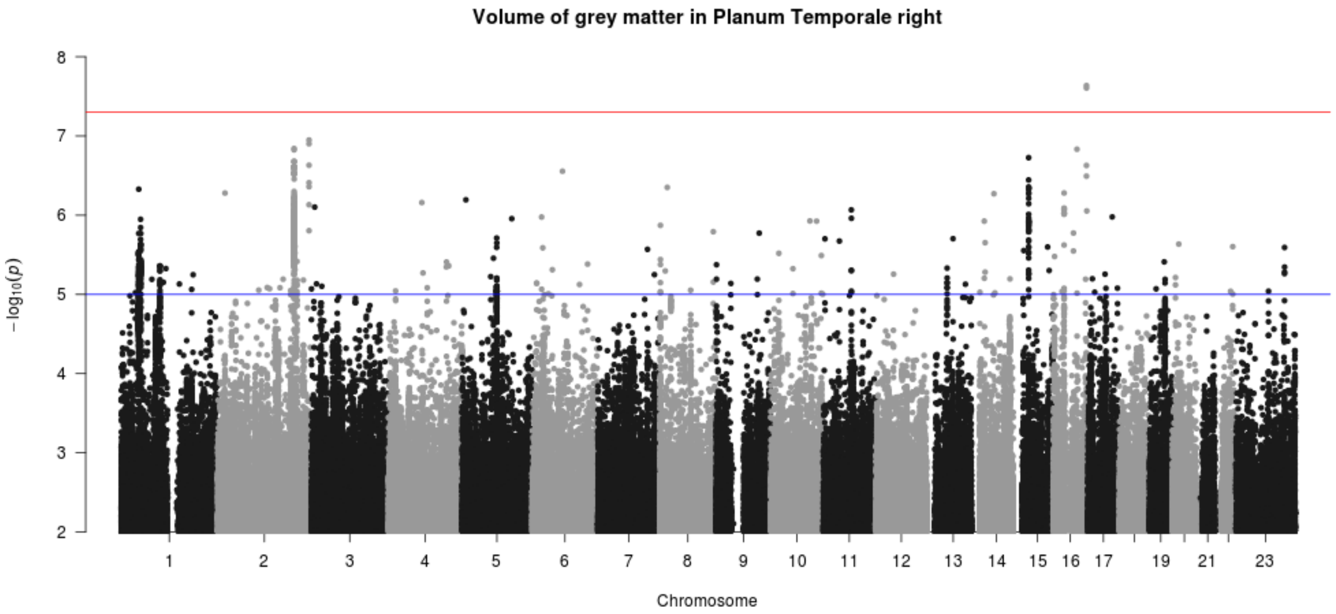


**Figure S6: QQ plot and Manhattan plot from the GWAS for right PT.**

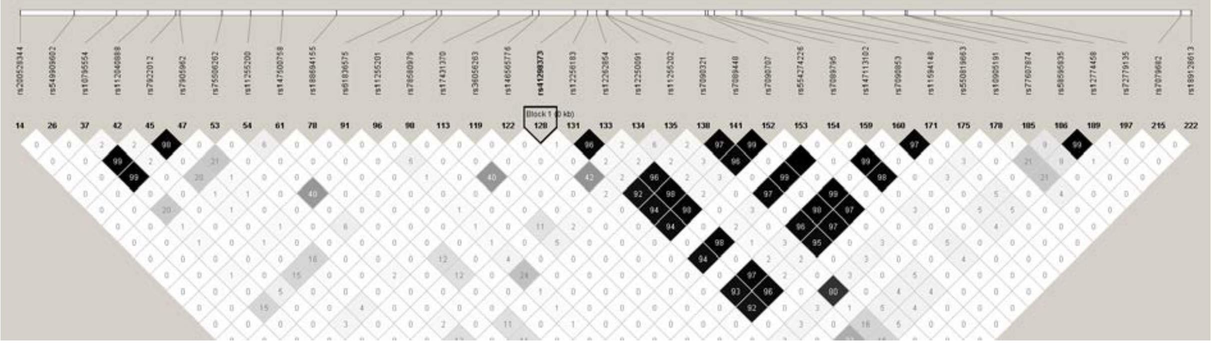

**Figure S7: Linkage disequilibrium ($r^2$) structure around rs41298373 (2kb flanking), calculated using the UKB imaging dataset and visualized using HaploView.**

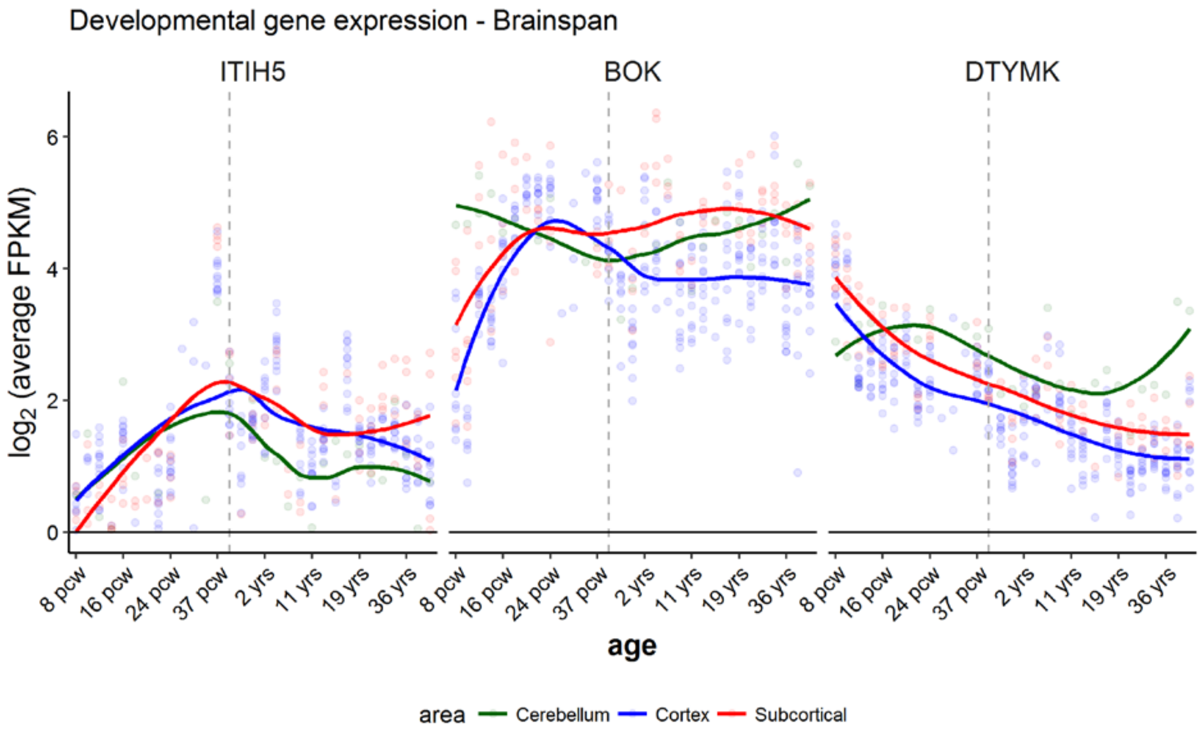


**Figure S8. Developmental expression of genes of interest which arose from the GWAS for PT asymmetry, according to the BrainSpan atlas.** Each dot indicates averaged expression (one to three independent measurements per gene, region, and time point). Colours indicate cerebellar, cortical or subcortical structures. Each dot shows the average expression within a brain structure and time-point. The lines indicate the smoothed function of the $\log_2$(average FPKM) across cortical, cerebellar or subcortical structures in relation to age.

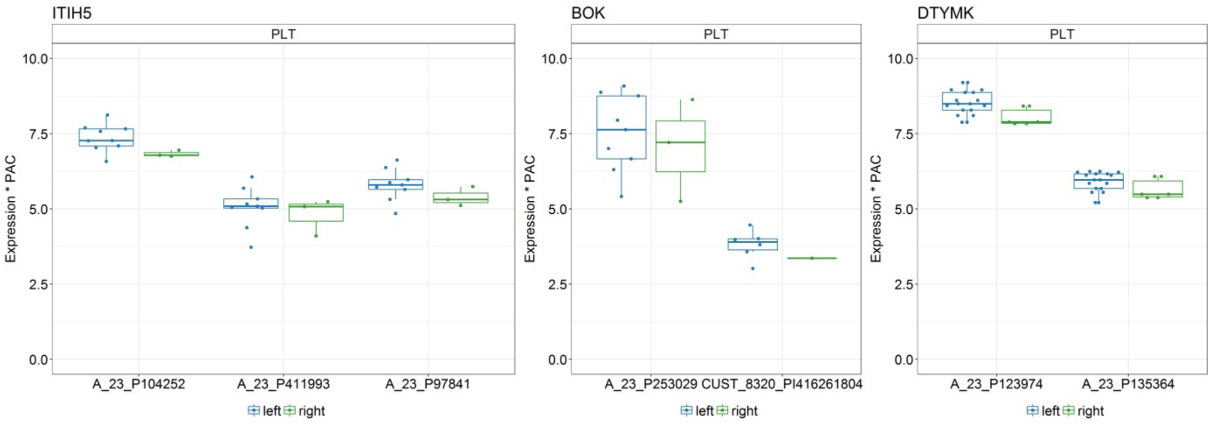


**Figure S9: Expressionof the genes of interest in the planum temporale region as defined by the Allen Brain Atlas.** The expression level is shown per microarray probe and hemisphere, for probes that were significantly different from the background (PAC=1).

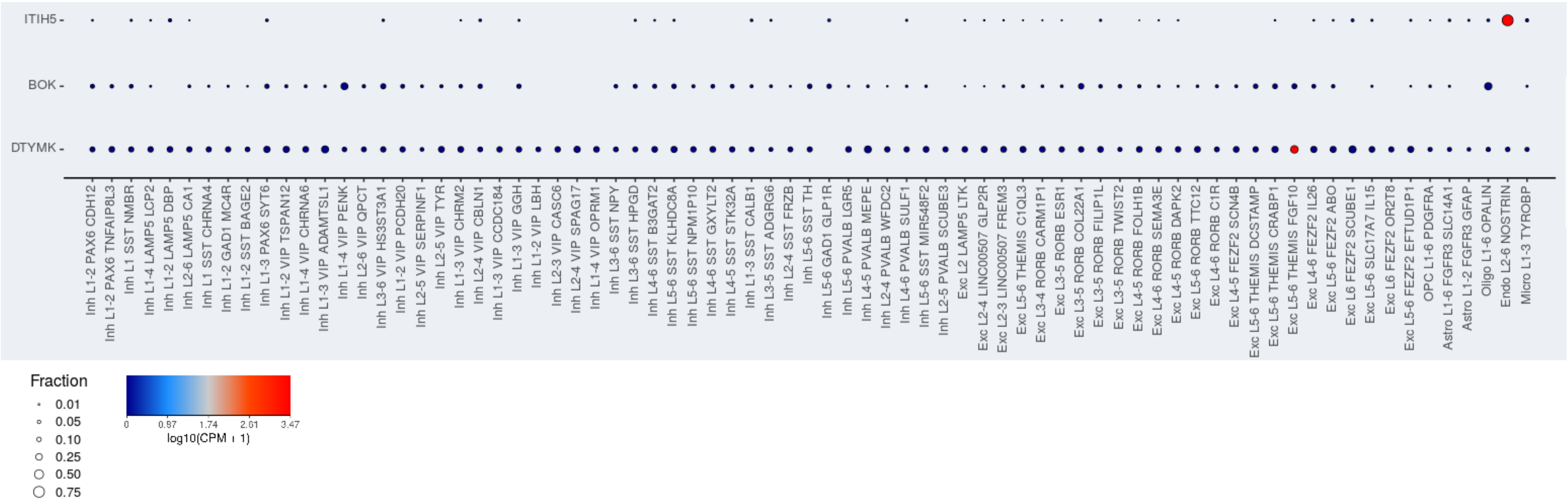


**Figure S10: Expression of the genes of interest across cell type clusters within human middle temporal gyrus (Allen brain atlas, http://celltypes.brain-map.org/rnaseq).**

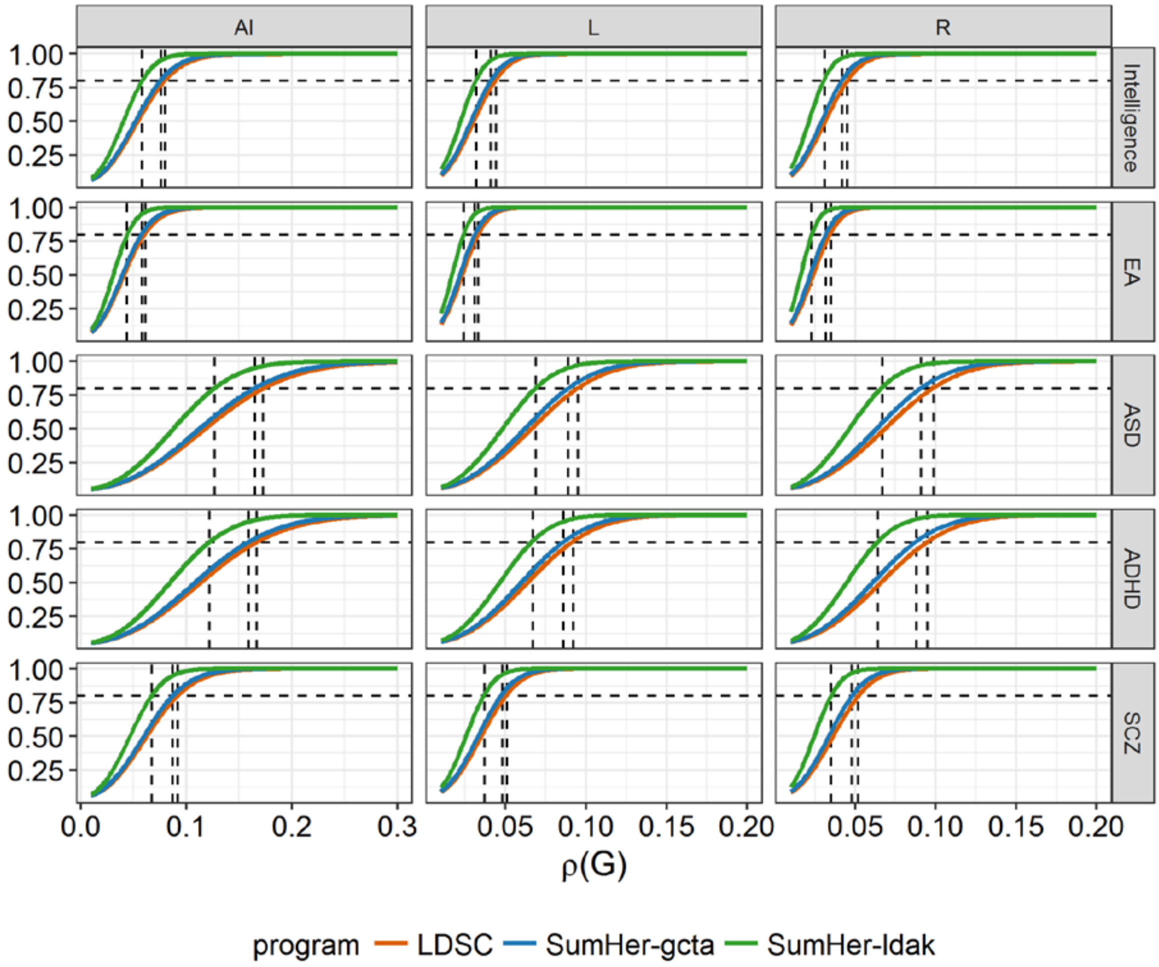


**Figure S11**: **Approximate power to detect genetic correlation (ρ) between PT volume measures (columns) and psychiatric, cognitive and behavioural traits of interest (rows).** The GCTA power calculator was used in all cases, into which were input different estimates of SNP-based heritability ($h^2_{SNP}$) that were produced by each different method for measuring genetic correlation (LDSC, SumHer-gcta, SumHer-ldak). See Figure S3 for the $h^2_{SNP}$ estimates for the PT measures, and Table S7 for the $h^2_{SNP}$ estimates for the other traits of interest. The dashed horizontal line indicates β=0.80. The vertical lines indicate the minimum genetic correlation (ρ value) detectable at β=0.80, assuming the method-specific SNP-based heritabilites.

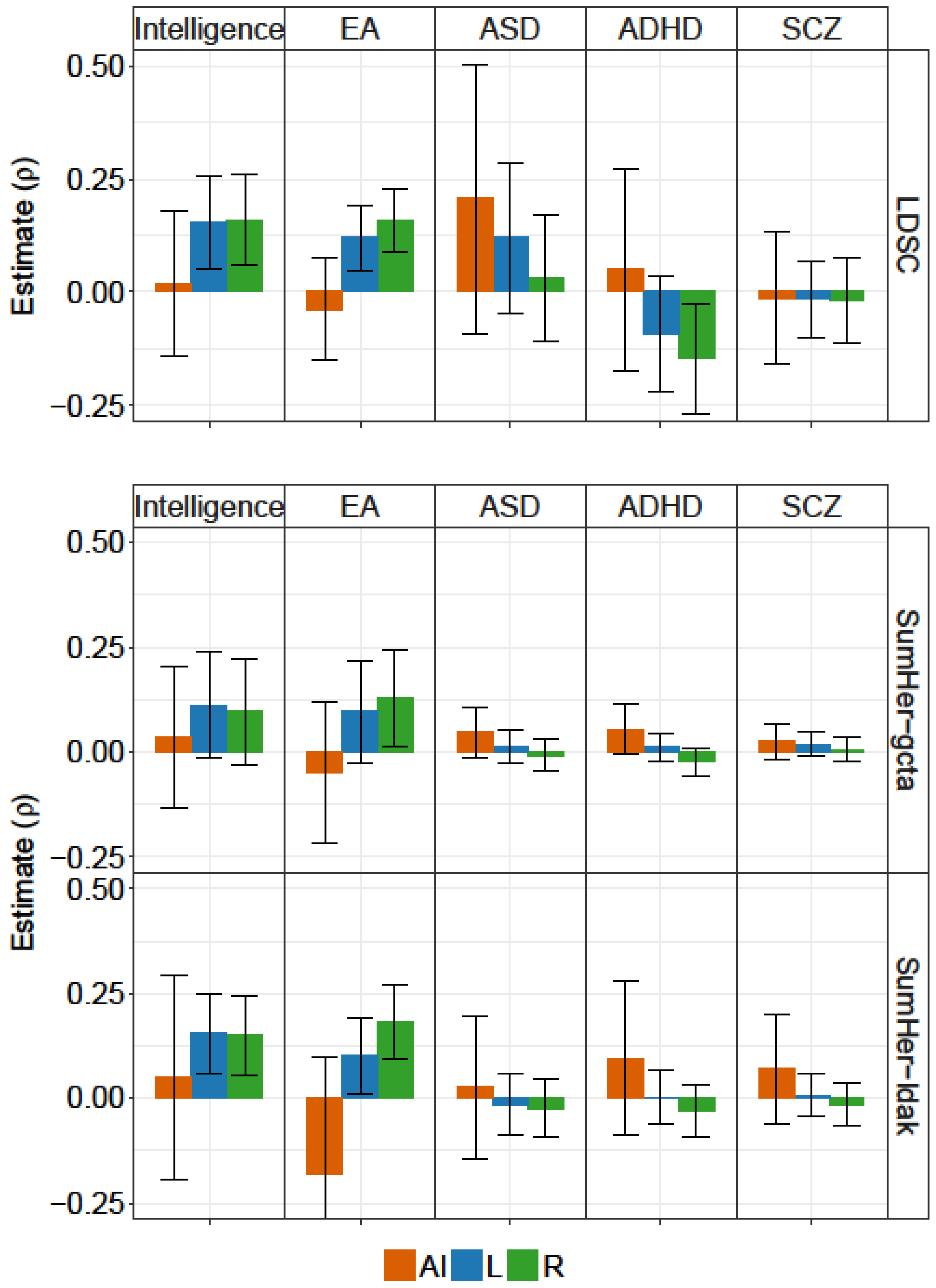


**Figure S12: Genetic correlations (ρ) between PT volume phenotypes and other traits of interest, estimated using LDSC and SumHer (gcta and ldak models).** The lines indicate the 95 % confidence intervals for the estimates.
